# Oxygen partitioning into biomolecular condensates is governed by protein density

**DOI:** 10.1101/2024.05.03.592328

**Authors:** Ankush Garg, Christopher Brasnett, Siewert J. Marrink, Klaus Koren, Magnus Kjaergaard

## Abstract

Biomolecular condensates form through the self-assembly of proteins and nucleic acids to create dynamic compartments in cells. By concentrating specific molecules, condensates establish distinct microenvironments that regulate biochemical reactions in time and space. Macromolecules and metabolites partition into condensates depending on their interactions with the macromolecular constituents, however, the partitioning of gases has not been explored. We investigated oxygen partitioning into condensates formed by intrinsically disordered repeat proteins with systematic sequence variations using microelectrodes and phosphorescence lifetime imaging microscopy (PLIM). Unlike other hydrophobic metabolites, oxygen is partially excluded from the condensate with partitioning constants more strongly modulated by changes in protein length than hydrophobicity. For repeat proteins, the dense phase protein concentration drops with chain length resulting in a looser condensate. We found that oxygen partitioning is anti-correlated with dense phase protein concentration. Several mechanisms could explain such an anti-correlation including excluded volume or salting out effects. Molecular dynamics simulations suggest that oxygen does not form strong and specific interactions with the scaffold and is dynamic on the nanosecond timescale. Biomolecular condensates thus result in variation of oxygen concentrations on nanometer length-scales, which may tune the oxygen concentration available for biochemical reactions within the cell.

## Introduction

Oxygen gradients form throughout biology when the rate of consumption exceeds the rate at which it can be replenished by diffusion. Oxygen gradients are well documented on meter scales in the ocean due to the respiration of organic matter, or on centimeter scales in e.g. lake bottoms (1–3). In our blood circulation, oxygen gradients occur on sub-millimeter scales across alveoli and arteries, where oxygenated blood releases oxygen to tissues (4). Diffusion works more efficiently at short length scales, and therefore it is increasingly difficult for oxygen gradients to arise. However, it has been speculated that oxygen gradients could also arise on a sub-cellular length scale, but this has been difficult to test experimentally. Nonetheless, oxygen gradients have been reported in giant umbrella cells, where deoxygenated regions formed around mitochondria due aerobic respiration (5). Similarly, swimming bacteria are capable of detecting changes in oxygen on µm scale of their long axis (6). Here we investigate an additional mechanism whereby oxygen gradients might arise on a subcellular length scale: passive partitioning into membrane-less organelles.

Cells are compartmentalized into several compartments. In addition to the traditional organelles, a new type of compartments has recently been identified that form by condensation of biomolecules, so-called biomolecular condensates, or membrane-less organelles (7,8). Like traditional organelles, membrane-less organelles can regulate biochemical reactions by spatial confinement – some components are concentrated, and others are excluded (9). Unlike traditional organelles, membrane-less ones can form and dissolve and thus serve as a trigger for changes in cellular metabolism or signaling (10,11). Oxygen is the ultimate oxidizing agent in most of our metabolism and thus takes part in many biochemical reactions, and in undesired side reactions for others. To understand how membrane-less organelles affect oxygen-dependent cellular processes, it is thus necessary to understand how oxygen partitions into biomolecular condensates.

Biomolecular condensates concentrate or exclude molecules through chemical partition equilibria. Condensates have different interior milieu compared to the bulk cytosol that form solvation-like interactions with other molecules and thus act as a distinct solvent phase (12,13). Macromolecules like proteins and RNA can be concentrated by thousands of folds or excluded because the condensate forms a mesh with pores too small to accommodate molecules beyond a certain size. Comparatively little is known about the partitioning of small molecules into condensates as these cannot easily be tagged with a fluorophore. Using fluorescence microscopy to quantify partitioning, it has been shown that condensates up-concentrate a range of small molecules in vitro and in cells (14–16), and that the protein scaffold creates distinct chemical environments that concentrate specific molecules. These studies were, however, limited to the study of fluorophores and thus mostly aromatic compounds. Mass spectrometry approaches developed for metabolomics allow investigation of a broader range of metabolites and drug-like compounds (17,18). This led to the conclusion that small molecule partitioning did not depend on high affinity binding with scaffold protein or pore size, but rather depended on solvent properties of the condensates, principally hydrophobicity. Investigation of condensate metabolomes in cells further showed that especially phospholipids are enriched in condensates (17).

A key challenge for the study of the sub-cellular oxygen gradients has been a scarcity of methods to detect such gradients. Phosphorescence lifetime imaging microscopy (PLIM) is currently the oxygen imaging technique with the highest spatial resolution. Some transition metal dye molecules display oxygen sensitive phosphorescence lifetimes, which has been developed into a series of probe molecules used for PLIM. The presence of heavy metals in these phosphorescent probes allows the efficient intersystem crossing from singlet excited state to triplet excited state leading to phosphorescence (19,20). The phosphorescence of these metal complex dyes, called indicators, gets quenched by molecular oxygen by the process of energy transfer from the indicator to the oxygen. We speculated that these dyes could be used to measure oxygen gradients inside biomolecular condensates as well.

In the present work, we measured oxygen partitioning in the biomolecular condensates using PLIM and found that oxygen concentrations are generally decreased inside condensates, a finding corroborated by molecular dynamics (MD) simulations. Our research highlights that oxygen partitioning is dominated by the dense phase protein concentration rather than the chemical properties of the scaffold.

## Materials and methods

### Protein expression and purification

Genes encoding octapeptide repeat protein [Q_5,8_]-20, [Q_5,8_]-30, [Q_5,8_]-40, [Q_5,8_]-60, [(3Q_5:_V_5_),Q_8_]-60, [(Q_5:_V_5_),Q_8_]-60 and [V_5,_Q_8_]-60 were synthesized and then cloned between the NdeI and BamHI sites in a pET-15b vector by Genscript (Table S1). Plasmids were transformed into chemical competent *E. coli* BL21(DE3) and grown at 37 ℃ in terrific broth containing 100 µg/mL ampicillin and 180 rpm shaking till stationary phase, where protein expression was induced with 0.5 mM IPTG and incubated for next 14-16 hours. Cells were harvested by centrifugation at 4000 g for 30 mins.

Repeat proteins were purified as reported with minor modifications (21). Briefly, cell pellet was dissolved in milli-Q water and lysed by probe sonication and pelleted by centrifugation (13,000 g, 30 min). The pellet was dissolved in a buffer (pH 7.4) containing 8 M urea+20 mM K_2_HPO_4_/KH_2_PO_4_, and 150 mM NaCl. Cell debris was cleared by centrifugation. The protein was purified by Ni-NTA affinity chromatography under denaturing conditions followed by 60 hours of dialysis. The protein solution turned turbid after dialysis and the pure protein was collected by centrifugation and lyophilized. The lyophilized repeat protein was reconstituted in 150 mM PBS (20 mM K_2_HPO_4_/KH_2_PO_4_, and 150 mM NaCl) buffer pH 7.4 heated at 80 ℃ for 10 minutes to reach a clear solution, which turned turbid when the temperature was decreased. This heated clear solution was quickly spun down to remove irreversible aggregates, if present. Protein concentrations were determined by diluting the sample until it was clear, measuring tyrosine absorbance at 280 nm and back calculating total concentrations using molar extinction coefficients from ProtParam and dilution factors. The isoelectric point of all the repeat proteins varies from 8.69 to 8.25 (from [Q_5,8_]-20 to [Q_5,8_]-60) as calculated using the ProtParam.

### Characterization of condensates

Condensates were imaged using a Zeiss Axio Observer Z1 inverted microscope where 40 μL of the turbid sample was drop-casted on the 10 mm microwell of 35 mm petridish (P35G-1.5-10-C from MatTek). The upper critical saturation temperature was determined by measuring the change in the optical density at 600 nm from 15 ℃ to 90 ℃ using a labbot (Probation Labs Sweden AB) UV-visible spectrophotometer. A homogeneous sample was heated at 90 ℃ which was cooled down stepwise till a final temperature of 15 ℃. The saturation concentration (C_sat_) was measured for all proteins by pelleting the dense phase by centrifugation follow by measurement of A_280nm_ in the dilute phase using the Labbot instrument (Labbot, Probation Labs Sweden AB).

### Dense phase concentration determination

The dense phase concentration of the droplets was determined by centrifuging phase separated samples at 13,000 g for 10 mins at a set temperature to get the continuous dense and dilute phase. A known volume of dense phase was taken out using positive displacement pipette and diluted in denaturation buffer (20 mM K_2_HPO_4_/KH_2_PO_4_,150 mM NaCl, 8 M urea) is required to get a homogeneous solution, followed by concentration measurement by UV-Visible spectroscopy at 280 nm.

### Electrochemical sensing of oxygen

Oxygen concentrations in the dense and dilute phases were measured using in-house built oxygen microsensor of diameter 10 µm also available at UNISENSE (Aarhus, Denmark) (22). These sensors measure oxygen reduction at a gold cathode and have a very low oxygen consumption between 0.4-5 nmol O_2_/hour. The calibrated (2 points in O2 free and air-saturated water) sensor was mounted on micromanipulator (MM33, UNISENSE) controlled by SensorTrace Suite via a motor controller (MOTCON). A macroscopically phase separated sample was prepared by centrifuging 100 µM of [Q_5,8_]-30 solution using swinging bucket centrifuge at 4000 g, and 20°C temperature for 10 mins. The sensor was inserted into the sample and moved across the phase boundary using the micromanipulator resulting in measurements of the oxygen concentration at different depths of the solution starting from several micrometers above the phase boundary.

### Phosphorescence lifetime imaging microscopy (PLIM)

Samples for PLIM experiments were prepared by adding a polar positively charged 500 μM dichlorotris (1,10-phenanthroline) ruthenium (II) hydrate (Sigma Aldrich) to condensate solutions (23). The concentration of the condensate sample used was just above the critical concentration to avoid spinodal decomposition and allow the formation of droplets in nucleation metastable region. Freshly made condensate samples were used to avoid sample variation due to aging of the condensates.

PLIM was performed using a Zeiss observer Z1 inverted microscope equipped with pco. FLIM lasers and frequency domain based luminescence lifetime camera system (24). The frequency modulation was set at 200 kHz based on the known unquenched lifetime of around 1 μs for the ruthenium dye. Starna fluorescent red (UMM/SFR/200X) has similar excitation wavelength as the ruthenium dye used and was used as a reference with a known lifetime of 5.25 ns. The laser exposure time was adjusted to 1 sec to get good intensity signals. The phosphorescent intensity of the ruthenium dye used as an indicator in the sample should be less than the intensity of the reference dye sample used to avoid background noise in the experiment. Oxygen concentrations were measured using a firestring needle type oxygen sensor with tip diameter of 430 µm (Pyroscience, GmbH), mentioned as reference oxygen sensor in the text, in the dilute phase of the samples. The oxygen sensor was calibrated using two known oxygen concentrations, air saturated water and an oxygen free solution (alkaline ascorbate solution).

We used a chemical oxygen scavenger i.e., sodium sulphite to create different oxygen concentration environment in the samples. We incubated the sample with different concentrations of sodium sulphite and measured the oxygen concentration in the dilute phase of the same sample using the reference oxygen sensor, where 10 mM sodium sulphite resulted in anoxic environment in the dilute phase. Further, we recorded the phosphorescence lifetime values in the dense phase of the same sample using PLIM to determine the partitioning coefficient from the Stern-Volmer plot. Further the oxygen concentration in the dense phase (i.e., inside the condensates) was extrapolated from the partitioning coefficient values obtained. PLIM images were analyzed using image J (Fiji software): Background noise was reduced using a Gaussian filter (sigma radius of 70), and the filtered image was subtracted from the original raw intensity image to reduce the background noise. The resulting intensity image obtained was redirected to lifetime channel to measure the phosphorescence lifetime of the dye inside the droplets. We analyzed the droplets by filtering for circularity and diameter and obtained the average lifetime and size for individual droplets. The mean phosphorescence lifetime of the dye in the droplets was plotted versus area of the droplets for both the anoxic and oxic conditions. Finally, we averaged the phosphorescence lifetime values for different sizes of the droplets and plotted the average phosphorescence lifetime of the dye inside the condensates against the known oxygen concentration in the dilute phase.

### Partitioning of fluorescent dyes into condensates

We monitored the partitioning of ruthenium dye (used in PLIM experiments) at a concentration of 500 µM for [Q_5,8_]-X repeats, where X ranged from 20 to 60, as well as for hydrophobic variants of [Q_5,8_]-60 using confocal microscopy (Zeiss LSM 90). Similarly, we have measured the partitioning of another three small-molecule fluorescent dyes—Sypro Orange, Fluorescein, and Rhodamine B—into [Q_5,8_]-20 and [Q_5,8_]-60 condensates. Fluorescein and Rhodamine B were used at concentrations of 20 µM, while Sypro-Orange was used at a 10 X concentration. Images were captured using a 63X objective lens. We performed Z-stacking to cover the region of entire droplet in a plane and generated 2D projections with maximum intensity to capture the complete fluorescence intensity within the droplets. The images were analyzed using Fiji software. Three images were taken for each set. Histograms were generated for each image and fitted with a Gaussian distribution to obtain the mean intensity and standard error of mean (SEM) of the dilute phase. For droplet identification, 2D projections were thresholded using a dark background and Otsu’s algorithm. The mean intensity and their corresponding SEM of droplets was calculated, with circularity parameters set between 0.2 and 1. The ratio of mean fluorescence intensities and their SEM inside and outside the condensates was used to calculate the partition coefficient of the fluorophores within the condensates.

### MD Simulations

Topologies and initial conformations for Martini 3 models of the IDPs were generated using Polyply, including updated bonded parameters for disordered proteins (25,26). Additionally, non-bonded protein-water interactions were increased by a factor of 1.04 to both improve the global dimensions of disordered proteins and reduce protein-protein interactions to avoid aggregates (27,28). A factor of 1.04 was used instead of the generally recommended 1.10 as this degree of increase was found to produce homogeneous systems with no protein phase separation. As described elsewhere, the increase in the interaction strength was achieved using a virtual site built directly on top of the protein backbone bead with a specific interaction to water (29). Molecular oxygen was represented by a single TC3 bead, based on water-octanol partitioning along with preliminary studies of membrane partitioning and binding to folded proteins (26,30). The parameters for HEME were taken from the work of Chiariello et al (31)

To simulate a system representative of a protein in the dilute region of a condensate system, a single protein was placed in a 20 nm cubic box, solvated, neutralized, and salt was added to a concentration of 150 mM NaCl. Oxygen beads were further added to the system at a concentration of 150 mM. For simulations of [Q_5,8_]-20 condensates, a slab configuration was prepared with a condensate of 50 molecules contained within the central 20 nm of a 12 x 12 x 60 nm simulation box. As mentioned before, the system was similarly solvated, neutralized, and ions and oxygen beads were both added to concentrations of 150 mM.

All simulations were performed using Gromacs 2021.7 (32). Systems were energy minimized, followed by equilibration and production simulations. Systems were equilibrated for 100 ns using a timestep of 10 fs and using the Berendsen barostat and thermostat (33). For production runs, a timestep of 20 fs was used for simulations of length 10 μs, using a velocity-rescaling thermostat and Parrinello-Rahman barostat (34,35). For single molecule simulations, the pressure coupling was maintained isotropically at 1 bar, while for slab condensate simulations, semi-isotropic pressure coupling was applied. All simulations were maintained at a temperature of 300 K. Non-bonded simulation parameters otherwise followed the new-rf Martini parameter set, with the exception of the new advised treatment of neighbor lists for the correct calculation of pressure, such that parameters of rlist =1.35 nm and verlet-buffer-tolerance = −1 were used (36,37). All simulations were performed with three replicates to ensure convergence and consistency in results.

Simulations were analyzed using a combination of Gromacs tools (gmx density) and custom analysis using Python scripts utilizing MD Analysis (38,39). Analysis was performed across the final 2 μs of the simulation time. Radial distribution functions were measured between protein backbone beads and oxygen molecules. To measure contacts between oxygen beads and proteins inside the condensate, a contact was defined as an oxygen being within 6 Å of any protein bead. The length of an individual contact event was defined as the number of consecutive simulation frames where an individual oxygen molecule was in contact with a protein. Similarly, the length of a single condensate visit was defined as the length of time a single oxygen was inside the bulk region of the condensate before leaving again.

## Results

### Synthetic IDP as model system for determining oxygen concentration in condensates

We aimed to study how oxygen partitions into biomolecular condensates (**Fig. 1a**), and thus needed a model system that is both representative of natural condensates and experimentally convenient. Many model intrinsically disordered proteins phase separate in the presence of crowding agents like PEG. However, these crowders also affect oxygen solubility (40), which is undesirable in this study. Therefore, we used a synthetic intrinsically disordered protein model system consisting of octapeptide repeats that form condensates at low saturation concentrations (C_sat_) without a crowding agent (**Fig. 1b**) (21). The repeat protein also allows systematic variation of the number of repeats and the properties of the sequence by changing the amino acid composition. The repeat protein i.e., GRGDSPYS originally contained serine in positions 5 and 8, which we replaced for glutamine, i.e., GRGDQPYQ for compatibility with other experiments. The repeat protein is charge neutral but has a slight positive charge coming from the termini. In the following, we refer to these as [Q_5,8_]-X, where X indicates the number of repeats.

**Figure 1:**
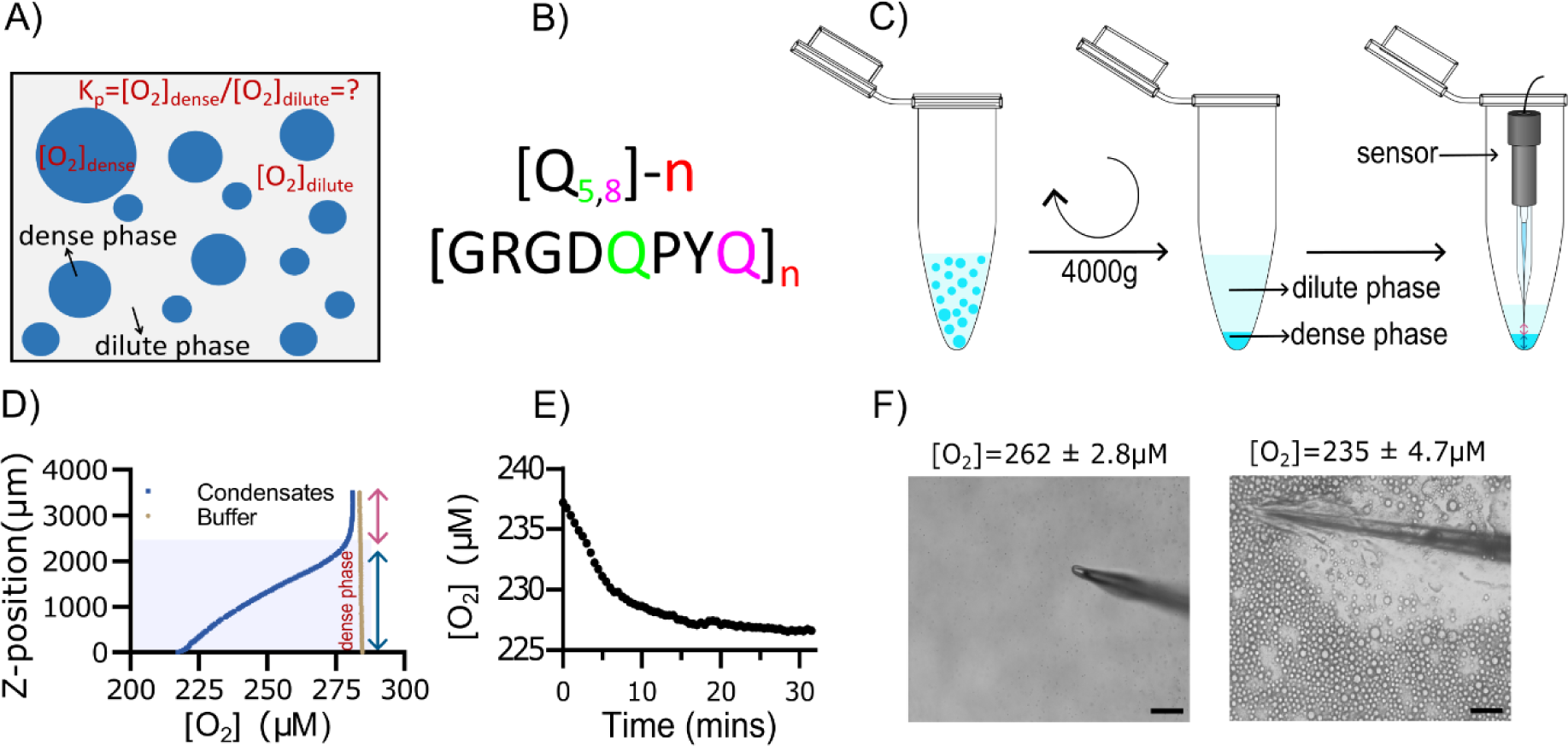
Determination of oxygen concentration in condensates using electrochemical sensing. (A) Oxygen concentrations in the dense and dilute phases are related by the partitioning coefficient (K_p_). (B) Artificial IDPs based on octa-peptide repeats form condensates and serve as a model system for oxygen partitioning. (C) A macroscopically condensate was prepared by centrifugation 4000 g of a phase separated sample of [Q_5,8_]-30 and Oxygen concentrations were measured across the phase boundary by inserting an electrochemical microelectrode into the continuous dense phase of [Q_5,8_]-30. (D) Change in the oxygen concentration (µM) with change in the sensor position from dilute phase to the dense phase (µm), where shaded blue area corresponds approximately to the pelleted [Q_5,8_]-30 condensate. (E) Time dependent change in the oxygen concentrations inside the continuous dense phase at a known depth. (F) Measurement of oxygen inside the biphasic system with the help of optical microscope with 20X objective using oxygen microelectrode where left panel indicates dilute phase and right panel indicates dense phase. Scale bar shown is 100 µm

Initially, we expressed and purified [Q_5,8_]-30.When reconstituted in PBS, these proteins form micrometer-sized spherical assemblies, which fuse upon encounter and grow with incubation time (**Fig. S1a**). The droplets are polydisperse with most having a diameter in the range 2-12 µm (**Fig. S1b**). These peptide repeats follow upper critical solution temperature (UCST) which means they form a two-phase system at lower temperature and turn into single homogeneous phase above a critical temperature (**Fig. S1c**). In agreement with previous characterization (21), this suggests that [Q_5,8_]-30 exhibits prototypical UCST liquid-liquid phase separation and is thus a good model for biomolecular condensates formed by many IDPs.

### Estimation of oxygen partitioning using a microelectrode

To measure oxygen concentrations inside biomolecular condensates, we initially used an electrochemical Clark-type microsensor with a tip diameter of 10 μm. We created a macroscopically phase separated sample by centrifugation, which resulted in a dilute phase on top of a protein rich dense phase (**Fig. 1c**). The oxygen profile was measured as the sensor was inserted across the phase boundary (**Fig. 1d**). The oxygen concentration was constant in the dilute phase, but gradually dropped with increasing depth, whereas no change was seen for buffer. The slope of the oxygen concentration dropped with increasing depth, suggesting an approach to a plateau. The electrochemical measurements indicated that the homogeneous dense phase partially excludes oxygen. The gradient observed by the microelectrode spans more than two millimeters, which is a much broader transition than expected from partitioning across a phase boundary. To test whether this represents a true gradient, we measured the oxygen concentration at a static depth over half an hour and observed a gradual approach to steady-state (**Fig. 1e**). The oxygen-consumption of the electrode is minuscule and is too low to cause a gradual depletion of oxygen, so instead we hypothesized that the gradual transition represented the interaction between the electrode and the condensates driving a deformation of the condensate. To test this, we measured oxygen concentrations in a phase separated sample with dispersed droplets, while observing the tip of the electrode under a microscope (**Fig. 1f**). We observed that the droplets collapse onto the electrode merging into film-like condensate on the tip of the electrode. This is similar to the wetting of surface observed for many condensates. Simultaneous measurement of oxygen concentrations showed an oxygen concentration between dilute solution and the steady-state value inside the condensate. The electrochemical measurements show that the oxygen concentrations can be measured inside macroscopically phase separated samples and the oxygen is depleted inside the [Q_5,8_]-30 condensates. However, the sensor tip perturbs the phase-separated sample, and it is not likely to be able to probe oxygen distributions in dispersed condensates. Therefore, we sought to develop a non-perturbing technique that measures oxygen concentration in co-existing phases.

### Measurement of oxygen inside the condensates using phosphorescence lifetime imaging

Phosphorescence lifetime imaging microscopy (PLIM) has been used to quantify the oxygen concentration in for example mammalian cells and organoids (41–45). The phosphorescent dye is quenched by the triplet state of molecular oxygen and the lifetime thus decreases with increasing oxygen concentration (**Fig. 2A**). We speculated that PLIM would also be useful for measuring oxygen partitioning in phase separated systems. As a sensor, we used dichlorotris (1,10-phenanthroline) ruthenium (II) hydrate (**Fig. 2A**), which is a water soluble positively charged dye, has excitation and emission in the visible range and has been used for PLIM oxygen imaging. In the following, we will refer to it as the ruthenium dye. We incubated the ruthenium dye with a phase separated sample of 20 μM [Q_5,8_]-30, and we imaged the condensates at 40X magnification using PLIM to record the phosphorescence intensity. Initially, we compared total dye concentrations from 100 μM to 1 mM (**Fig. S2**) aiming to find the highest concentration where lifetimes are not affected by self-quenching. Phosphorescence lifetimes are similar for 100 μM and 500 μM, whereas 1 mM dye results in shorter lifetimes suggesting undesirable self-quenching. We thus used a total 500 μM dye in all subsequent PLIM experiments to maximize signal-to-noise.

**Figure 2:**
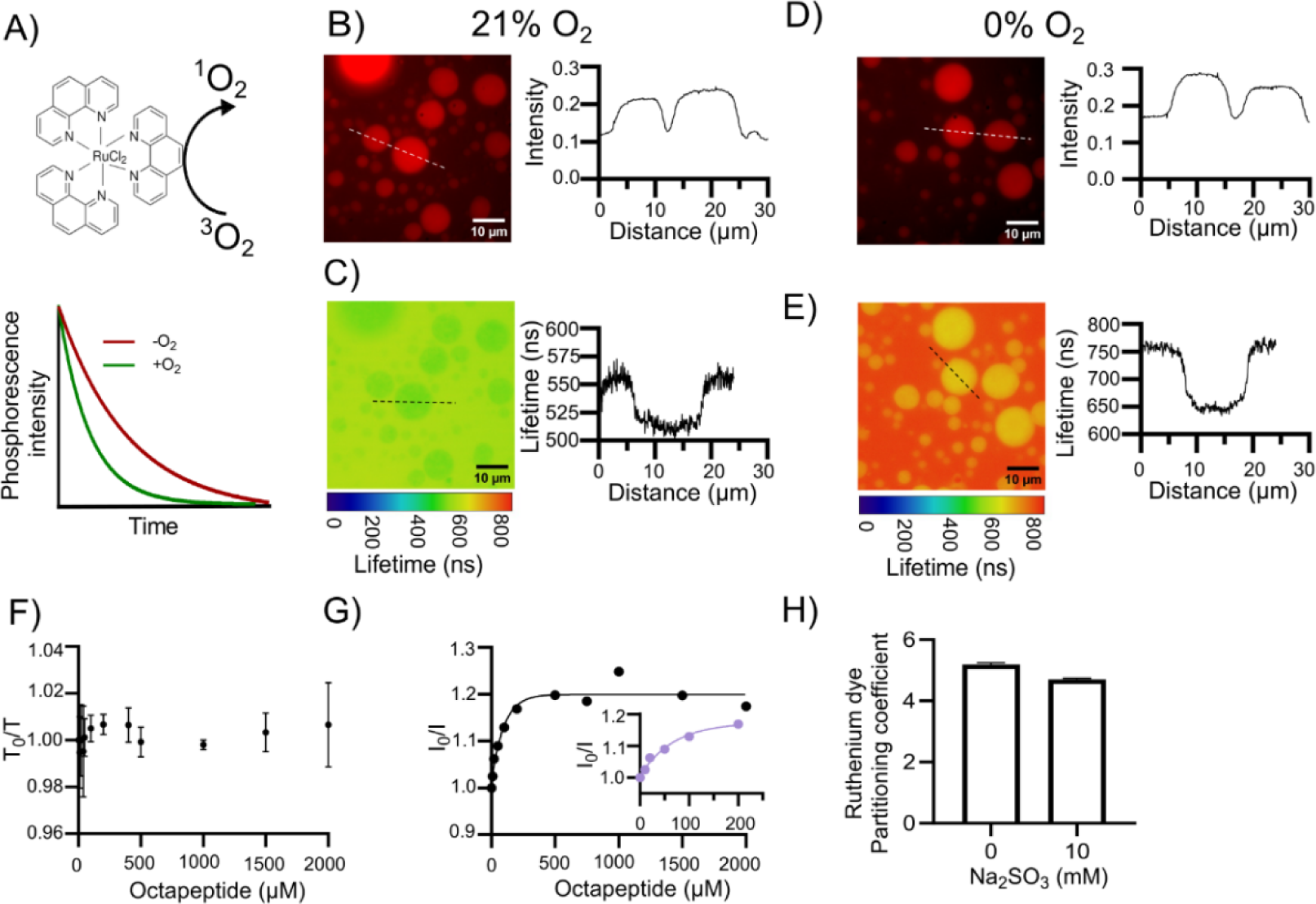
Determination of oxygen concentrations in condensates using PLIM. (A) Phosphorescence from ruthenium-dyes is sensitive to the oxygen concentration and the excited stated is quenched by contact with triplet state oxygen. Phosphorescence lifetimes thus decrease in the presence of oxygen, which allows oxygen concentrations to be imaged using PLIM. The chemical structure represents the dye used throughout this study named Tris dichlorotris (1,10-phenanthroline) ruthenium (II) hydrate where (B, D) represented Phosphorescence intensity and (C, E) corresponded for lifetime images of a phase separated sample of [Q_5,8_]-30. (B, C) were recorded on samples in equilibrium atmospheric oxygen concentrations, and (D, E) under anoxic conditions. Intensity and lifetime line profiles were extracted at the indicated dashed lines. (F) Time resolved fluorescence studies showed that phosphorescent lifetime of the ruthenium dye is similar at all concentrations of the peptide used. (G) Stern volmer plot of the interaction between ruthenium dye and peptide indicates weak interaction between the dye and the peptide. (H) The partitioning coefficient of the ruthenium dye into the condensates is similar in the presence and absence of sodium sulphite (Na_2_SO_3_), used to create anoxic environment.

Condensates can be distinguished as round objects of elevated intensity (**Fig. 2B**) and changed lifetimes (**Fig. 2C**), which allows determination of the lifetime in the dense and dilute phases simultaneously. Cross-sections of droplets reveal a steep gradient in the phosphorescence lifetime between a high value in the dilute phase and a lower value in the dense phase (**Fig. 2C**). In principle, the gradient could represent a boundary layer zone between the phases. However, the spatial resolution of the imaging system is likely not sufficient to resolve the boundary layer, so we focus on the lifetime in the center of the condensate. To quantify oxygen concentrations, we measured the phosphorescence lifetime in the absence of oxygen by adding sodium sulphite, a chemical oxygen scavenger (**Fig. 2(D-E)).** Neither sodium sulfite nor the dye affects the phase behavior of the condensates at the concentrations used as assessed by the critical concentration and temperature of phase separation (**Fig. S3**). Removal of oxygen increased phosphorescence lifetimes in both the dense and the dilute phases but remained lower inside the condensates (**Fig. 2E**). This suggests that the dye experiences a different environment inside the condensate which changes its phosphorescence emission. It is necessary to account for this change in basal lifetime when quantifying oxygen concentrations.

The change in phosphorescence lifetime could either be due to an interaction between the dye and the IDP, or an emergent property of the dense phase. To distinguish these possibilities, we used an octapeptide which has the same sequence-repeat as the condensate forming IDPs but does not undergo phase separation. We titrated the dye with the octapeptide while recording phosphorescence lifetime and intensity (**Fig.**). Addition of the peptide did not affect the phosphorescence lifetime but led to a ∼20% reduction of phosphorescence intensity. The curve shows clear saturation in the high-micromolar range indicating a weak, but specific interaction with the peptide resulting in slight static quenching. These interactions also lead to preferential partitioning of the dye into the condensates shown by a partition coefficient of 5.19±0.065 as measured by comparing intensity by confocal fluorescence microscopy (**Fig. 2H**), which is similar to the partitioning of other hydrophobic small molecules into condensates (14,18). In total, this suggests that the change in lifetime is not due to a simple collisional quenching by the condensate protein but is an emergent property of the different physical environment inside the condensate that has to be accounted for.

Oxygen concentration can normally be determined from single lifetime measurement, when the lifetime can be compared to reference lifetime measurements at known oxygen concentrations – for example recorded via an electrode. This assumes that the unquenched lifetime does not change between the measurement and the reference which is not the case for the PLIM measurements in condensates. To determine the oxygen concentrations in the dense phase, we thus need to take the changed lifetime into consideration although we cannot get direct reference values from electrode measurements. Instead, we referenced used the dilute phase as a reference, where matching values of τ, τ_0_ and K_SV_ can be obtained.

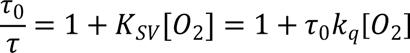

This allows us to determine k_q_, which is determined by how often collisions between molecules result in quenching and is likely to be similar between the two phases. This allows us to determine the oxygen concentration inside condensates from values of τ and τ_0._

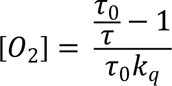

For measurements of dense phase lifetime as a function of oxygen concentration, the lifetimes have to be plotted against [O_2_]_dilute_, in which case K_P_ can be determined from the slope:

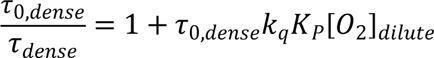

To quantify the oxygen concentration, we measured phosphorescence lifetimes in [Q_5,8_]-30 condensates at different sodium sulphite concentrations that only partially scavenge the dissolved oxygen (**Fig. 3A**). Oxygen concentrations were estimated by a titration in buffer sample which was observed over time following addition of sodium sulphite (**Fig. S4**). 10 mM Na_2_SO_3_ resulted in complete removal of oxygen after three minutes, whereas lower concentrations gradually decreased to a near steady-state plateau. We performed PLIM imaging 25 minutes after addition of Na_2_SO_3_, which allowed us to estimate the oxygen concentration from the standard curve. Phosphorescence lifetimes decrease with increasing oxygen concentration as expected from a collisional quenching mechanism (**Fig. 3B**) resulting in a straight line in a Stern-Volmer (**Fig. 3C**).

**Figure 3:**
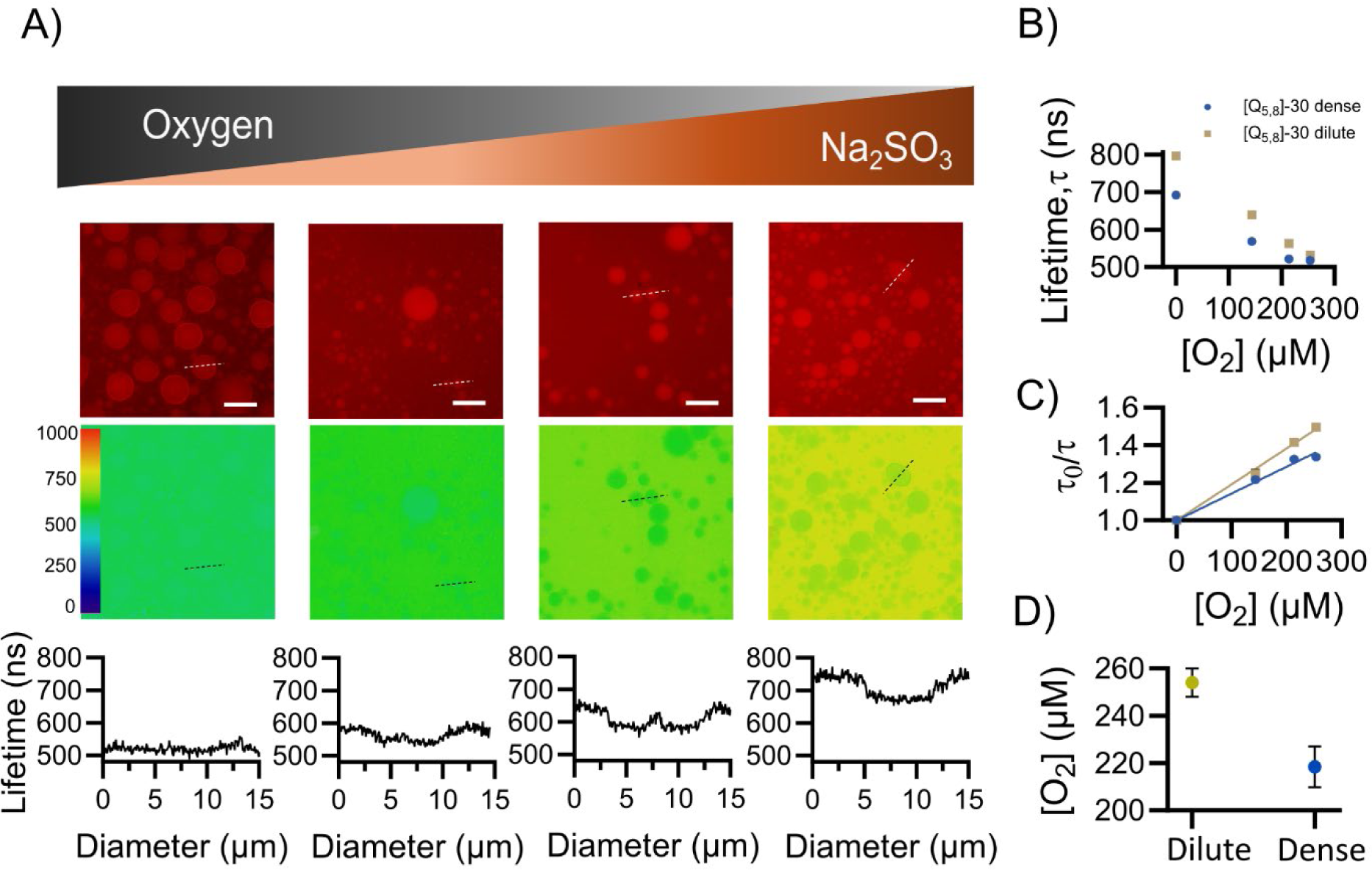
Oxygen dependent phosphorescence lifetime in the droplets. (A) Phosphorescence lifetime imaging of the [Q_5,8_]-30 condensate sample at different oxygen concentrations created by created by titrating with sodium sulfite (Na_2_SO_3_), where phosphorescence intensity is shown in top panel, phosphorescence lifetime in middle panel and line profile of the lifetime across droplet in bottom panel, where panel 1-4 represent increasing Na_2_SO_4_ from 0mM, 0.5mM, 1mM to 10mM corresponding to oxygen concentration of 254 µM, 214 µM, 144 µM to 0 µM respectively (B) Plot of Phosphorescence lifetime versus known oxygen concentrations determined using optode measurement (C) Stern Volmer plot where τ_0_/τ plotted with change in oxygen concentration for dense phase droplet (blue) and dilute phase (brown) of the [Q_5,8_]-30 condensates. (D) Oxygen concentrations in the dense and dilute phases as determined from the Stern-Volmer plot. Error bars represent standard deviation (S.D.) from triplicate readings.

K_P_ obtained from the slope was approximately 0.86 ± 0.027 and the oxygen concentration in the dilute phase was 254 µM ± 6.0 µM (**Fig. 3D**). Therefore, the [O_2_]_dense_ was 218 µM ± 8.6 µM (**Fig. 3D**). In agreement with microelectrode measurements, this indicated that protein condensates partially exclude oxygen and create low oxygen zones inside them. While it is possible to measure oxygen lifetimes at intermediate oxygen concentrations using chemical oxygen scavengers, the oxygen concentration is only stable at atmospheric and completely anoxic conditions. In the remaining experiments, we thus focus on these two conditions, which still allows us to determine oxygen concentrations as described above.

### Determinants of K_p_

Next, we set out to investigate the determinants of oxygen partitioning by varying the condensate properties. Condensates are polydisperse in nature with large size distribution, so first we checked the effect of the size of the droplets on the phosphorescence lifetime. Phosphorescence lifetime decreases slightly for bigger droplets (**Fig. S5**), but condensates at atmospheric oxygen and anoxic conditions were affected similarly, so the difference effect mostly cancels out for estimations of oxygen concentrations.

Oxygen is non-polar with an octanol-water partition coefficient of 4.4 (46), which suggested that it is more prone to partition into hydrophobic condensates in line with other small molecules (14,18). Therefore, we wanted to modulate the hydrophobicity without disrupting phase separation. Therefore, we started from a longer repeat protein [Q_5,8_]-60, which forms condensates at very low protein concentration, and substitute glutamine at the 5^th^ position to valine. To get a range of hydrophobicity, we substituted every fourth repeat [(3Q_5_:V_5_),Q_8_]-60, every other repeat [(Q_5_:V_5_),Q_8_]-60 or every repeat [V_5_,Q_8_]-60. Proteins were purified and characterized by microscopy and UV-visible spectroscopy (**Fig. S6**). All variants produced round condensates, however [V_5_Q_8_]-60 mostly formed irreversible aggregates, and the droplets formed were below a diameter of 2 µm. The samples are heated at 80 °C temperature to solubilize the condensates and quickly spin down at high speed i.e., 14000g to remove irreversible aggregates. However, the aggregation propensity indicated that we cannot further increase the hydrophobicity of the peptide repeats. The critical concentration gradually decreased with increasing hydrophobicity **(Fig. S6B).** Partitioning coefficients of the dye were within error for the different condensates tested (**Fig. S7**). We observed decrease in droplet size with increasing hydrophobicity. To avoid confounders, we restricted our analysis to droplets with diameters from 1 to 4 μm that were present in all variants.

Next, we measured the oxygen concentration in the dilute homogeneous solution of [Q_5,8_]-60, [(3Q_5_:V_5_),Q_8_]-60, [(Q_5_:V_5_),Q_8_]-60 and [V_5_,Q_8_]-60 using a firestring oxygen optode as a control for PLIM experiment. We observed that the oxygen concentration in the dilute phase of all hydrophobic variants is similar around 265 µM ± 10 µM. We measured the phosphorescence lifetime of [Q_5,8_]-60, [(3Q_5_:V_5_),Q_8_]-60, [(Q_5_:V_5_),Q_8_]-60 and [V_5_,Q_8_]-60 condensates at atmospheric oxygen condition and anoxic conditions to determine the oxygen concentrations in the dense phase. The difference in phosphorescence lifetime from oxic-to-anoxic condition decreased with an increasing hydrophobicity for similar size of the droplets with exception of [V_5:_Q_8_]-60 (**Fig. 4A-D**). Next, we averaged the phosphorescence lifetime for all droplets in the similar size distribution range and plotted the ratio of phosphorescence lifetime in anoxic to oxic condition (τ_0_/τ) i.e., against dilute phase oxygen concentration to calculate partitioning coefficients. Unexpectedly, oxygen showed a non-monotonous dependence on condensate hydrophobicity. Oxygen partitioning into the condensates decreased upon substitution of every fourth glutamine but increased upon further substitutions (**Fig. 4E**). Substitutions of residues in the octa-repeat may also affect the condensate interior in other ways than via hydrophobicity. To probe the condensate structure, we measured the protein concentration inside the droplets. The dense phase concentration of [(3Q_5_:V_5_),Q_8_]-60 variant was similar to [Q_5,8_]-60 variant but decreases for [(Q_5_:V_5_),Q_8_]-60 variant. We could not experimentally determine the dense phase concentration for [V_5_,Q_8_]-60 due to very low volume fraction (**Fig. 4G**). The decrease in the oxygen concentration for [3Q_5_:V_5_),Q_8_]-60 with similar dense phase concentration could be the compaction of the condensates and decrease in solvent availability. However, there is no clear correlation between the dense phase concentration and oxygen partitioning observed for other hydrophobic variants due to the change in both the hydrophobicity and dense phase concentrations. Further, it is difficult to interpret the effect of hydrophobicity in these variants due to very low volume fraction of the condensates.

**Figure 4:**
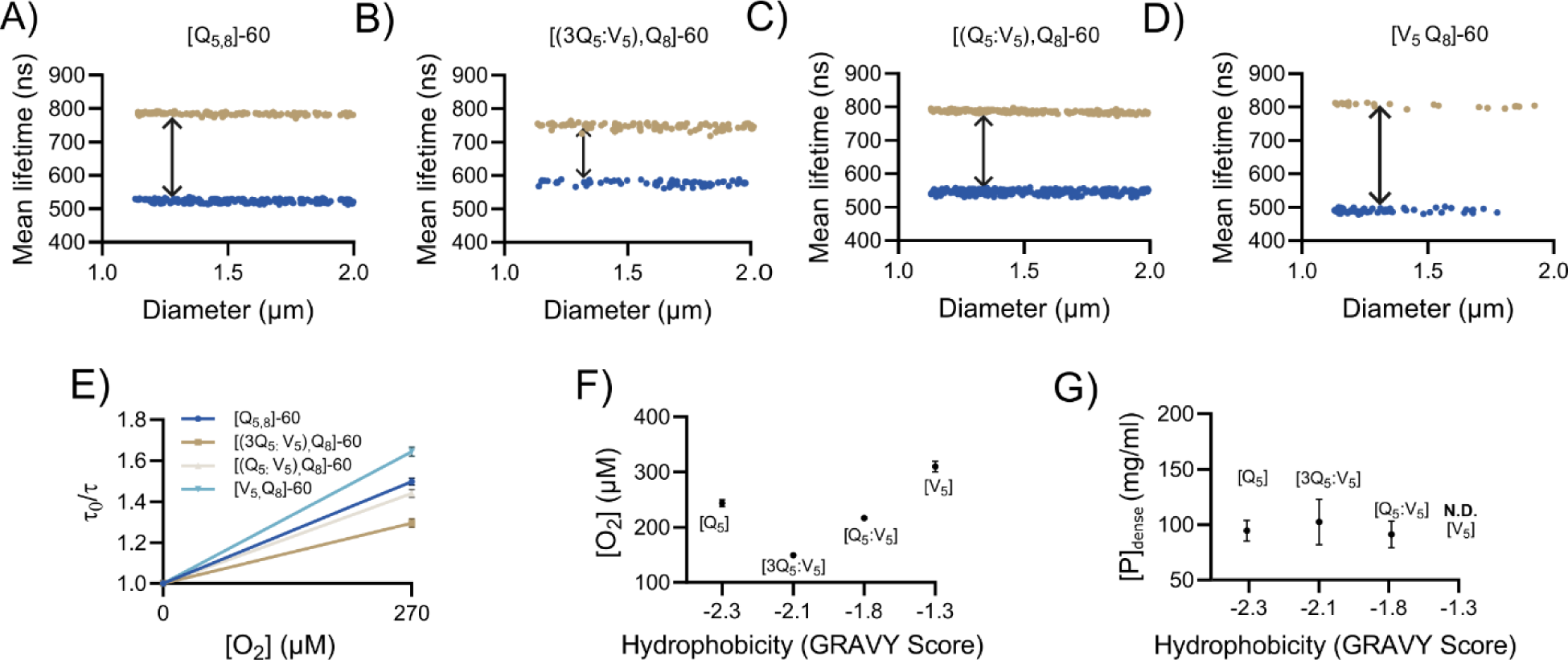
Effect of hydrophobicity on the partitioning of oxygen in the droplets. (A-D) Mean phosphorescence lifetime versus condensate diameter for condensates with different hydrophobicity. (E) Stern-Volmer plot for all the hydrophobic variants to estimate the effect of hydrophobicity on oxygen quenching. (F) Concentration of oxygen in the droplets was plotted with change in hydrophobicity (GRAVY score) number. (G) Dense phase concentration of different variants is plotted with change in hydrophobicity (GRAVY score), where the dense phase concentration for [V_5_] variant cannot be determined. Error bars represent standard deviation (S.D.) from triplicate readings.

To explore the determinants of oxygen partitioning further, we sought a way to change the phase diagram without changing the chemical environment in the condensates. Uniquely, repeat proteins offer this possibility as the phase diagram is highly sensitive to the number of identical repeats. We thus expressed and purified [Q_5,8_]-20, [Q_5,8_]-30, [Q_5,8_]-40 and [Q_5,8_]-60 and characterized them as above (**Fig. S8**). All the variants formed condensates with condensate size and critical concentration decreasing with chain length. The oxygen concentration as measured by the oxygen optode in the dilute phase of all variants is similar around 265 µM ± 10 µM. We measured the phosphorescence lifetime of the similar sized [Q_5,8_]-20, [Q_5,8_]-30, [Q_5,8_]-40 and [Q_5,8_]-60 condensates at atmospheric oxygen condition and anoxic conditions to determine the oxygen concentrations in the dense phase (**Fig. 5A**). The difference in the phosphorescence lifetime from oxic-to-anoxic condition increased with an increasing number of repeats for similar size of the droplets (**Fig. 5A**). The average phosphorescence lifetime values for dyes inside the droplets was analysed to calculate K_p_ and the oxygen concentration inside the condensates (**Fig. 5B**). The K_p_ increased monotonously towards one with increased chain length suggesting less exclusion of oxygen for the longest chains. (**Fig. 5C**). The effect was larger and more consistent than for variants with varied hydrophobicity. To exclude a technical origin of this effect, we repeated the measurements using continuous dense phases pelleted by centrifugation and insertions of an electrochemical sensor as in Fig. 1. The oxygen concentrations inside the dense phase of the condensates follow the same trend regardless of detection method (**Fig. 5C**).

**Figure 5:**
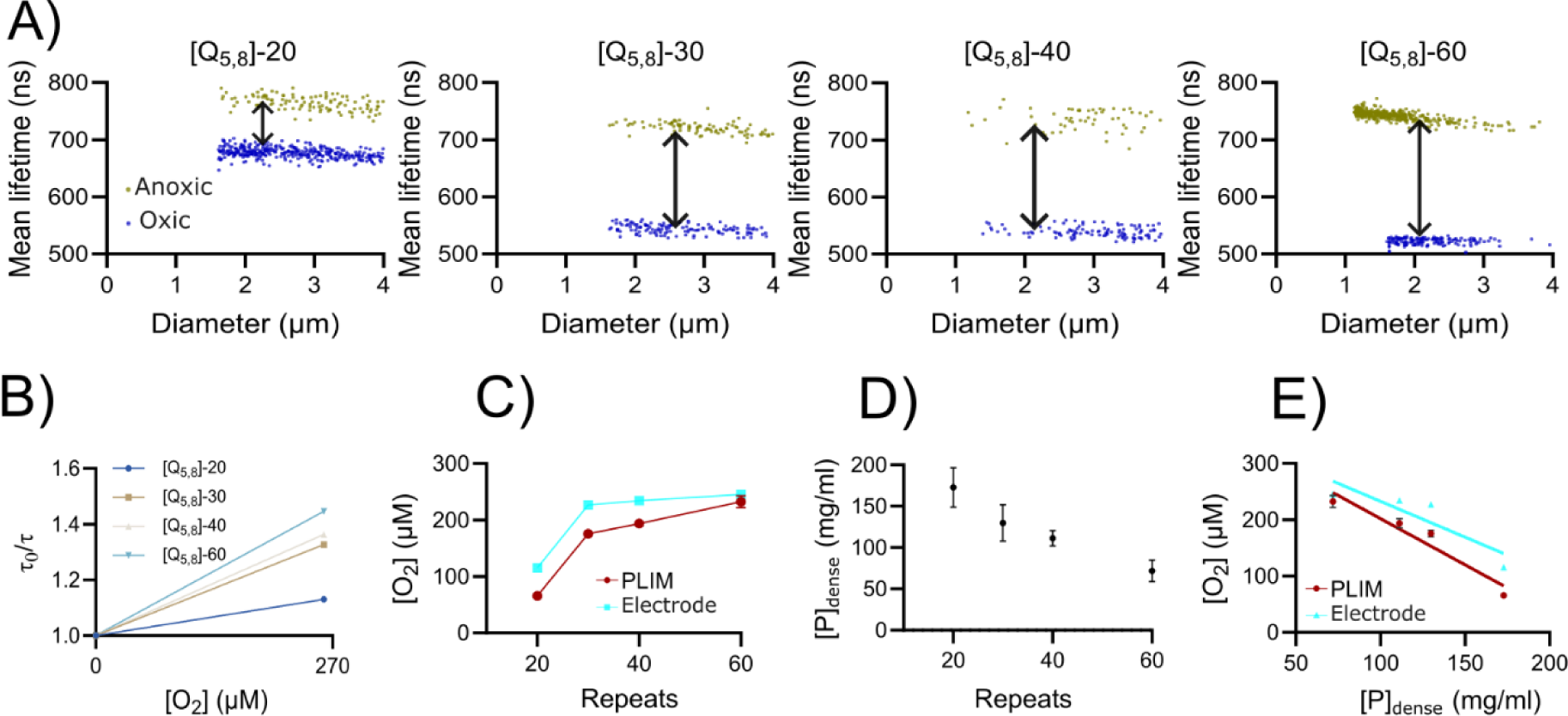
Effect of chain length on the partitioning of oxygen. Correlation between mean phosphorescence lifetime and droplet size for condensates formed by (A) [Q_5,8_]-20, [Q_5,8_]-30, [Q_5,8_]-40, and [Q_5,8_]-60. The difference between the lifetime under anoxic and oxic conditions increased with repeats as shown with double-sided black arrows. (B) Stern Volmer plot for all the repeats to estimate the effect of repeats on phosphorescence lifetime and partitioning coefficient determined from the slope of the curves. (C) Oxygen concentrations are recorded in the droplets using PLIM (red) and in the continuous dense phase with electrode (cyan) with change in the number of repeats. (D) Dense phase concentration of different protein variants is plotted with change in number of repeats and (E) Oxygen concentration is plotted with change in dense phase concentration using PLIM (red) and Electrode (cyan). Error bars represent standard deviation (S.D.) from triplicate readings. **Note**: The dense phase determination for the repeats (figure 6) and the hydrophobic variants (figure 5) has been done separately.

The internal structure of the condensates could also act as a potential determinant of partitioning. Condensates formed by different chain length are likely to have different internal organization although they appear similar under a microscope. We turned to dense phase protein concentration as an overall reporter of condensate organization. Chain length generally reduces the critical concentration of repeating polymers, but much less is known about how it affects the dense phase concentration. We measured the dense phase concentration for [Q_5,8_]-X and found that it decreased uniformly with chain length (**Fig. 5D**). [Q_5,8_]-20 has protein concentration of ∼170 mg/mL, which is similar to values previously reported for other condensates (48–51). For [Q_5,8_]-60, the dense phase protein concentration is much lower (∼70 mg/mL) suggesting that the long repeat proteins form a much looser condensate. Oxygen and protein concentrations in the dense phase were anti-correlated (**Fig. 5E**) with Pearson correlation coefficients of −0.96 (PLIM), −0.88 (electrode).

Both the exclusion of oxygen and the dependence on chain length was unexpected. To test whether this is a property unique to oxygen, we measured partitioning of three fluorophores in [Q_5,8_]-20 and [Q_5,8_]-60 using fluorescence microscopy. All three fluorophores partition favorably into both condensates, although to different extents with K_P_ varying from ∼2 to 8. The difference between [Q_5,8_]-20 and [Q_5,8_]-60 was small consistent with previous findings that partitioning is defined sequence properties and hydrophobicity (18). This suggests that the exclusion behavior of oxygen is not a shared by other hydrophobic small molecules.

### Molecular dynamics simulations of oxygen partitioning

To investigate how oxygen interacts with the condensates in atomistic detail, we thus undertook molecular dynamics (MD) simulation of oxygen partitioning. MD methods range from all-atom models that can represent any type of chemistry to highly coarse-grained models where each residue is represented by a single bead. Although it is possible to simulate at an all-atom resolution, these force fields are generally too computationally expensive to handle the system size needed to achieve phase separation (47). At a much coarser resolution, although one-bead-per-residue methods are efficient at capturing the phase behavior of IDPs, they are inherently unable to describe the interactions of small molecule clients such as oxygen (48–51). Between these options is the Martini force field, which uses a transferable building block approach representing 2-4 heavy atoms by a single bead, combining beads to build entire molecules, parameterized based on underlying chemical properties (26,52,53). Martini has already shown promise in simulating several condensate systems (28,54–56). Building on this, we showed that Martini resolution simulations can reproduce the phase separation of [Q_5,8_]-20 IDPs as expected (**Fig. S10**).

We added models of the dye and oxygen to condensate simulations to assess, how they interaction with the condensates. There is no Martini model for the dye used in this study, so instead we used HEME, which is also a large poly-aromatic ring system. HEME partitioned into the condensate as observed for the dye, but its presence did not affect the dynamics of the condensate as indicated by measurements of the protein diffusion coefficients through incoherent scattering functions (Fig. S11) (53). Qualitatively, the simulations agree with the experimental results in predicting an 50% lower oxygen concentration inside the condensate relative to the outside (**Fig. 6A**). Moreover, the visits of oxygen to the condensate region are typically very short lived. As **Fig. 6B** shows, single visits to condensate are predominantly less than 10 ns in total before the molecule leaves to the dilute region of the system again, with longer occupations becoming exponentially rare. This suggests that oxygen mostly remains highly dynamic inside the condensate.

**Figure 6.**
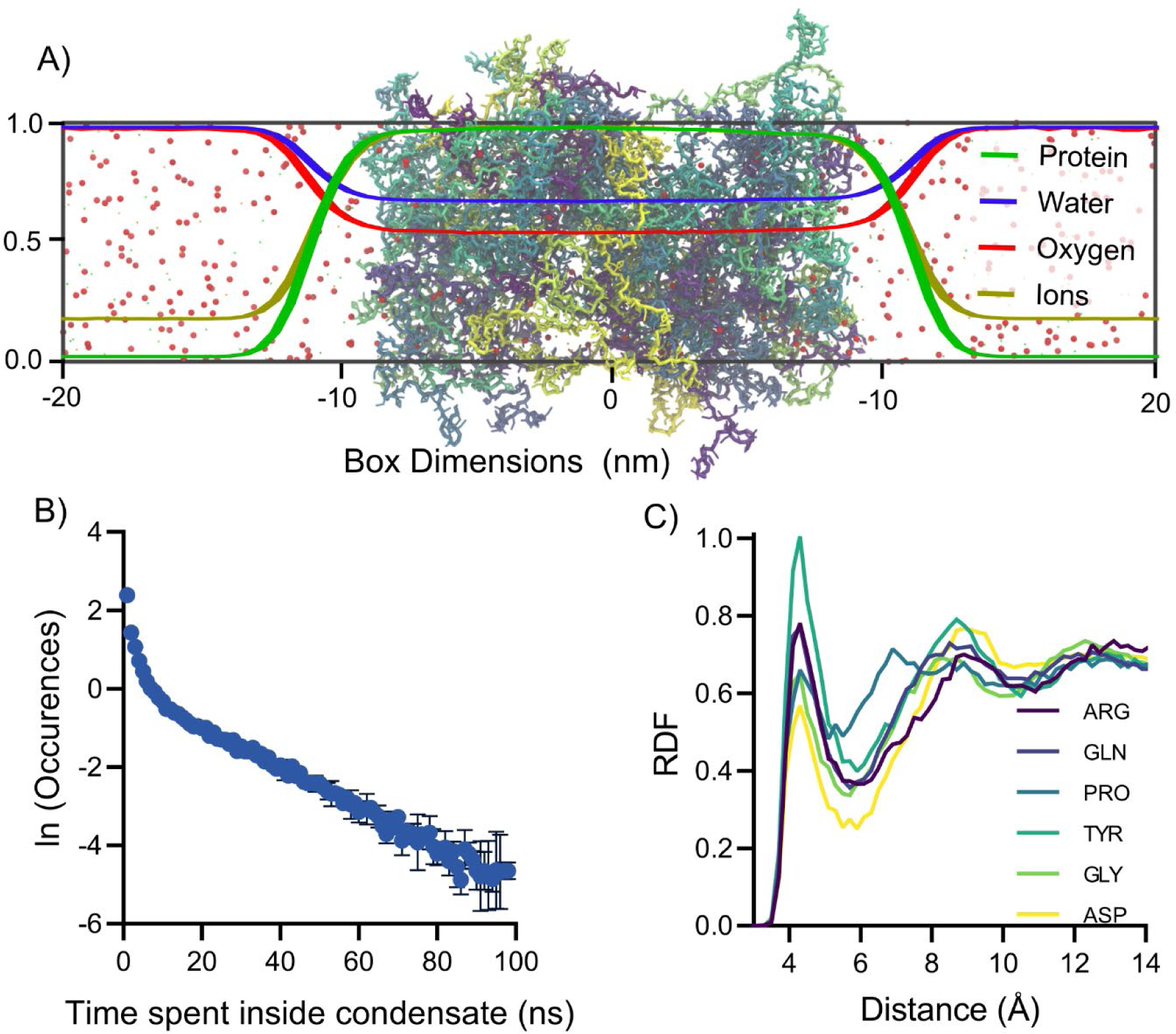
Coarse-grained simulations of oxygen molecules in condensates. (A) Normalized density profiles of the simulated system components across a slab condensate system. (B) Lengths of visits of oxygen molecules to the bulk region of the condensate. (C) Radial distribution functions plotted for the distribution of oxygen around different residues of the octapeptide repeat unit.

To understand the potential interaction between oxygen and the condensate forming proteins, we looked at the distribution of oxygen around each of the residues in the octapeptide repeat unit of the [Q_5,8_]-20 protein using radial distribution functions (**Fig. 6C, Fig. S12**). While there is relatively little change in the residue-specific radial distribution functions, there is a slight indication that oxygen is more prevalent around tyrosine compared to other residues in the octapeptide repeat. The same sequence of preference around the repeat unit residues is also observed for a simulation of a single protein (representing the dilute phase of a condensate system), further indicating that the overall interaction of oxygen with disordered proteins is weak and the condensate does not change the preference for interactions compared to the monomeric state (**Fig. S13)**. Biomolecular condensates can concentrate or deplete ions in their interior including the repeat proteins studied here (57). High ionic strength is known to reduce oxygen solubility (58) and could contribute to the exclusion of oxygen from the dense phase. To assess this effect, we analyzed the ion and oxygen concentrations in the absence of NaCl as well as at two different salt concentration. In agreement with previous studies, we find that NaCl is concentrated in the dense phase. Oxygen is still partially excluded from the condensate in the absence of NaCl, and increasing concentrations leads to slight decrease in oxygen partitioning. The absolute change is much smaller than the variation observed experimentally (**Fig. S14**). This suggests that salt partitioning is not the main mechanism of the exclusion of oxygen from condensates.

Finally, we investigated the hydrophobic sequence effect also investigated experimentally by titration in valine residues at position 5 in the octa-repeat. In simulations, the oxygen partitioning into the condensate increases monotonously with increased hydrophobicity (**Fig. S15**). The oxygen model is mainly parameterized based on e.g. water-octanol partitioning data, so that is the expected behavior of a hydrophobic solute. The experimental data show a non-monotonous trend with an overall larger magnitude of variation. The discrepancy suggests that the underlying cause is not well captured by the Martini forcefield yet.

## Discussion

In the present work, we developed and compared methods for measuring oxygen partitioning in biomolecular condensates. We have applied this technique to synthetic condensates composed of intrinsically disordered repeat proteins with systematic variations. We found that oxygen was partially excluded from the dense phase, and that the extent of this exclusion varied between condensates with an anticorrelation to protein density.

Optical measurements - and especially fluorescence microscopy - have become one of the main techniques to probe the properties of biomolecular condensates as they can resolve the signals from the dense and dilute phase. Optical measurements allow the interior to be characterized in terms of intensity, wavelength or lifetime. Here we apply PLIM techniques, that has been developed for imaging on much longer length scales, to biomolecular condensates. Compared to electrochemical sensing techniques, PLIM has the advantage of working in a dispersed solution and not perturbing the phase separated. We think the main disadvantages of the PLIM technique is its reliance on an assumption that the quenching constant k_q_ is unchanged between the phases, and the inconvenience of making measurements at intermediate oxygen concentrations. Most PLIM studies assume invariant k_q_, which cannot be independently measured, and it is a prerequisite for converting lifetimes into corresponding oxygen concentrations. Variations in k_q_ normally arise from binding of the dye molecule or slowed diffusion, which alters its rate of collision with oxygen. While condensates retard the diffusion of macromolecules, small molecules have comparable diffusion constants to dilute buffers (59). The dye molecule interacts weakly with the condensate, however, not enough to change the properties of the condensate. While we can thus not exclude a small effect on k_q_, we do not think this is likely to affect the data much as there is a good agreement between optical and electrochemical oxygen measurements. This suggests that bias would be shared between the two orthogonal techniques and likely represent the chemical potential of oxygen.

A second limitation is that we resort to measuring oxygen concentration from two points only, atmospheric and anoxic conditions. We have tested two methods for measuring at intermediate concentrations, an environmentally controlled imaging setup allowing us to control the oxygen concentration in the gas phase and partial chemical oxygen scavenging. Oxygen readings in a controlled atmosphere was found to be impractical due to the relatively long time needed for the system to come to equilibrium. This is incompatible with many studies of condensates as they mature during the equilibration process and there will be non-negligible evaporation due to the small volume of the sample. Partial oxygen scavenging was more successful (**Fig. S4**) and allow us to measure a concentration dependence of the phosphorescence lifetime. However, such systems are only in a semi-stable steady-state with an associated error in oxygen concentration even in calibration curves. Measurement at the two concentrations that can easily be controlled is thus a practical approach to allow correction for the different unquenched lifetime in the dense phase, but do not allow detection of e.g. unexpected effects to the curve shape.

Recently, a series of papers have studied partitioning of small molecules into condensates by measuring partitioning of large libraries of small molecules into a few condensates (14–16,18). Our study is different in only focusing on a single client but studying its partitioning into condensates with systematic changes in sequence. The former is useful for training machine learning models, the latter is more suited for studying sequence-function relationships in condensates. This led to the surprising conclusion of a dense phase dependence of partitioning for oxygen. In the project design, hydrophobicity and chain length were meant as orthogonal properties due to the repeat protein design. This is true at the level of individual molecules, where GRAVY scores are similar for proteins of different length. However, when it comes to condensate properties, such orthogonality may break down as the protein is more hydrophobic than the solvent, and a higher protein concentration may thus also translate into a higher mean hydrophobicity of the dense phase. This, however, does not translate into a stronger partitioning of the aromatic dyes tested (**Fig. S9)**.

The dense phase of a biomolecular condensate represents a challenge to sequence-function relationships as many condensate properties are linked, which makes it difficult to definitely assign a mechanism to the changes in oxygen partitioning observed. However, oxygen seems to behave differently from most other small non-polar molecules as it is depleted rather than concentrated inside condensates and that this correlate to protein density. Several mechanisms could explain such a correlation. At the most basic, the protein density occupies a certain volume in the dense phase as densest condensate we have studied is close to 20% protein mass. In addition to this comes an associated solvation sphere from which oxygen is likely also partially excluded. An excluded volume effect would suggest why oxygen partitioning correlates with protein density but is not the only possible explanation. Oxygen solubility decreases with increasing salt concentration and the condensate can both concentrate ions through partitioning and the protein brings a high density of charged residues itself, which naturally also scales with protein density. In the densest condensate, the concentration of charged side chains can be estimated to 0.44 M. NaCl concentrations in this range results in approximately 10% in oxygen solubility. While such effects may contribute, they seem unlikely to account for full ∼70% depletion observed even when accounting for concentrations of counterions. Considering also the effect of hydrophobicity discussed above, there are at least three plausible mechanisms by which dense phase concentration can affect oxygen partitioning without considering sequence specific effects. In conclusion, this suggests that partitioning of oxygen – and by extension other small molecules – into condensates is considerably more complex than equilibria involving simple hydrophobic solvents.

In conclusion, we have developed tools for studying oxygen partitioning in condensates and discovered a so far unappreciated factor in partitioning of small molecules. A natural extension of this work is to wonder whether similar effects modulate oxygen partitioning into condensates in a cellular context. This study solely considers reconstituted systems in vitro and in silico, and cannot provide direct proof for oxygen partitioning in a cell. However, the mechanisms described here are likely to be general, and a recent study suggested that in vitro partitioning correlates with partitioning in cell lysate, although the latter generally had a smaller magnitude (18). Future studies will reveal whether oxygen partitioning in condensates *in vitro* are used by nature to modulate oxygen-dependent physiological process.

## Acknowledgements

This work was supported by grants from the Villum Foundation (40662) to M.K., and a Novo Nordisk Foundation Interdisciplinary synergy grant (NNF20OC0063808) to M.K. and S.J.M, Grundfos Foundation and Carlsberg Foundation to K.K. Authors would like to thank Lars Borregaard Pedersen and Mette Hoffmann Asmussen for excellent technical support.

## Notes

### Competing Interest Statement

The authors have declared no competing interest.

### Summary of Updates

The updated pre-print contains new experiments including a more detailed characterization of the condensates, control experiments that support the interpretation and a rewritten discussion section that describes several interpretation of the underlying mechanism.

