## Supplemental information for "Oxygen partitioning into biomolecular condensates is governed by protein density"

This file contains 15 supporting figures and 1 supporting table.

**Supporting Figures**

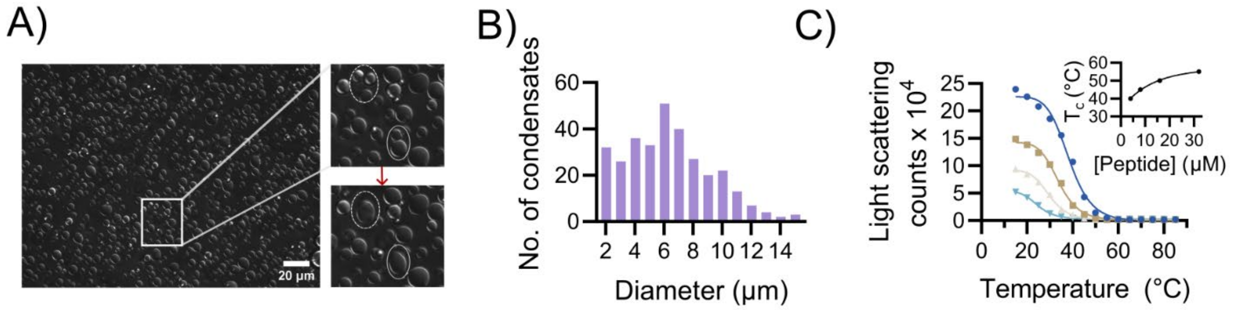


**Figure S1: Characterization of octa-repeat protein condensates.** (A) DIC microscopy images of [$Q_{5,8}$]-30 droplets. Enlarged view shows subsequent images in a time series showing fusion of droplets. (B) Size distribution of the [$Q_{5,8}$]-30 condensates from microscopic images using image J software. (C) Temperature dependent light scattering of [$Q_{5,8}$]-30 protein concentration of 4µM (light blue), 8 µM(beige), 16 µM (brown) and 32 µM (dark blue). (C) Inset shows protein concentration-dependent change in the critical temperature ($T_c$).

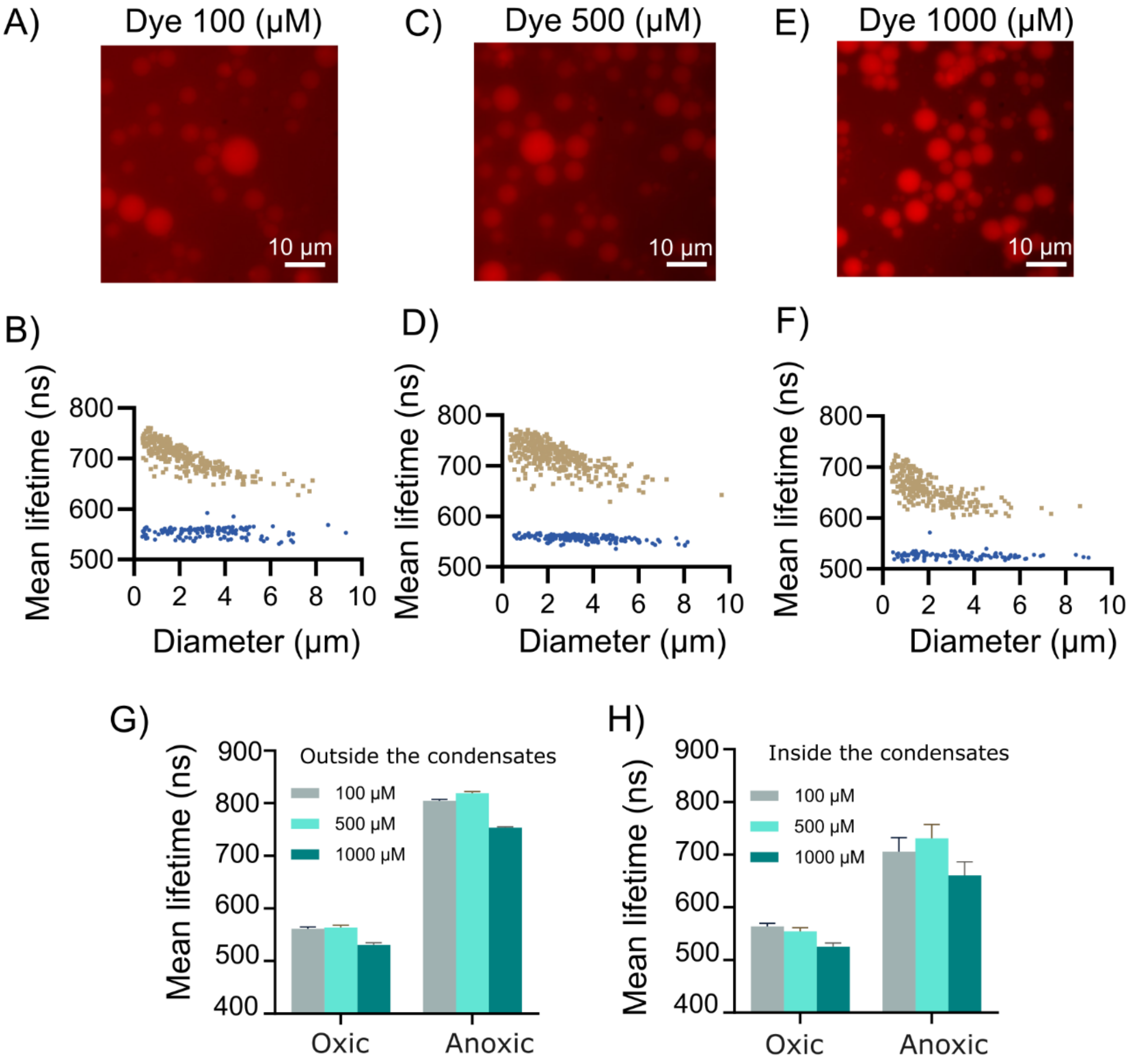


**Figure S2: Effect of total ruthenium dye concentration on lifetime measurements**. Microscopic images of the [$Q_{5,8}$]-30 condensates at 100 µM (A), 500 µM (C) and 1000 µM (E) dye concentrations. Mean lifetime of the phosphorescence dye inside the [$Q_{5,8}$]-30 droplets for anoxic condition (brown) and oxic condition (blue) for (B) 100 µM, (D) 500 µM and (F) 1000 µM of dye concentration. Mean lifetime of the ruthenium dye under oxic and Anoxic conditions at 100 µM (grey), 500 µM (light green) and 1000 µM (dark green) dye concentrations (G) outside the condensates and (H) inside the condensates.

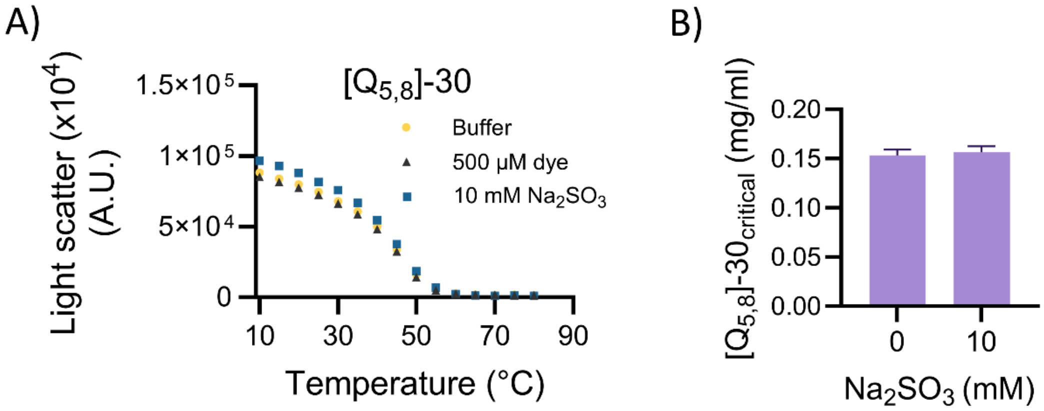


**Figure S3**: (A) Temperature dependent scatter experiment to undermine the effect of the sodium sulfite ($Na_2SO_3$) (10 mM) and the phosphorescent dye (500 µM) on the phase behavior of the [$Q_{5,8}$]-30 condensates. (B) The critical concentration i.e., the dilute phase concentration of the [Q5,8]-30 condensates in the presence and absence of 10 mM sodium sulfite ($Na_2SO_3$).

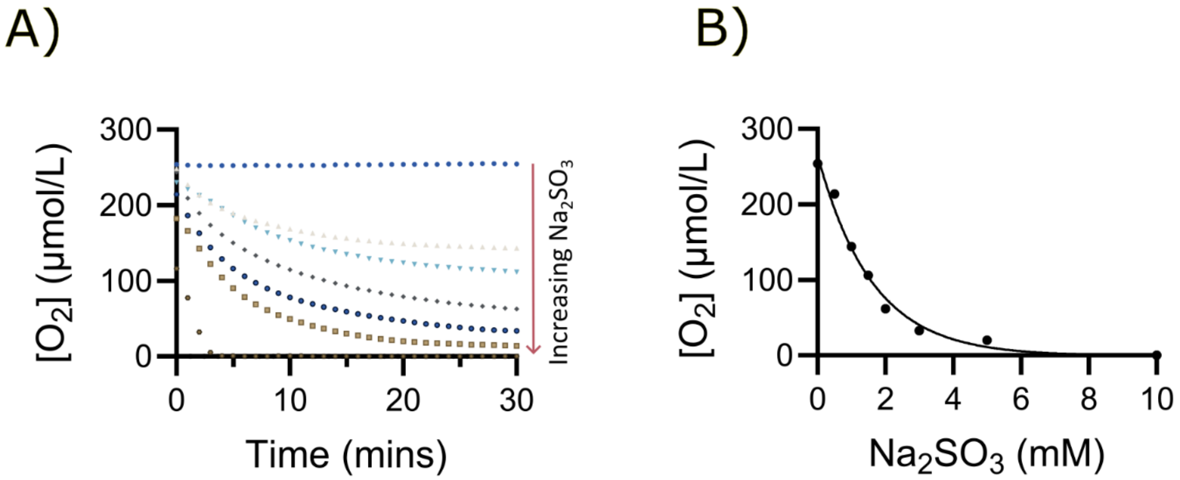


**Figure S4 :** Measurement of oxygen in the PBS buffer (20mM PB+150mM NaCl) at different concentrations of sodium sulfite using optode sensor. A) Time dependent change in the oxygen concentration at different concentrations of sodium sulfite used from 0mM to maximum 10mM concentration. B) Plot between the end point steady state oxygen concentration v/s sodium sulfite concentration.

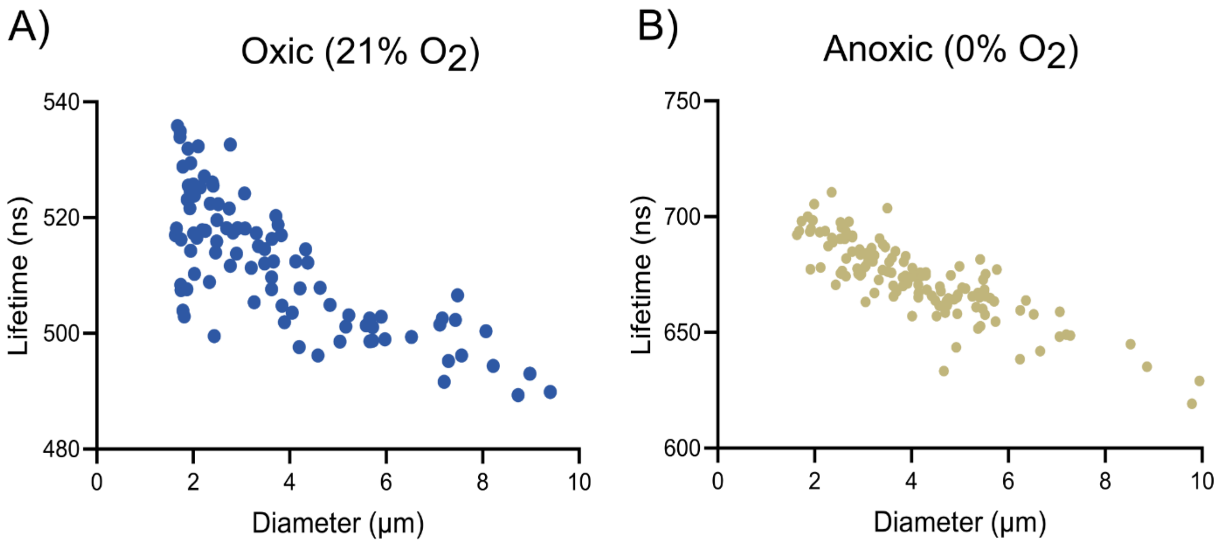


**Figure S5:** Effect of the droplet size on the lifetime of the dye inside the droplets. Lifetime of the dye inside droplets v/s cross sectional area of the droplets under (A) Oxic conditions i.e., 21% oxygen and under (B) Anoxic conditions i.e., 0% oxygen.

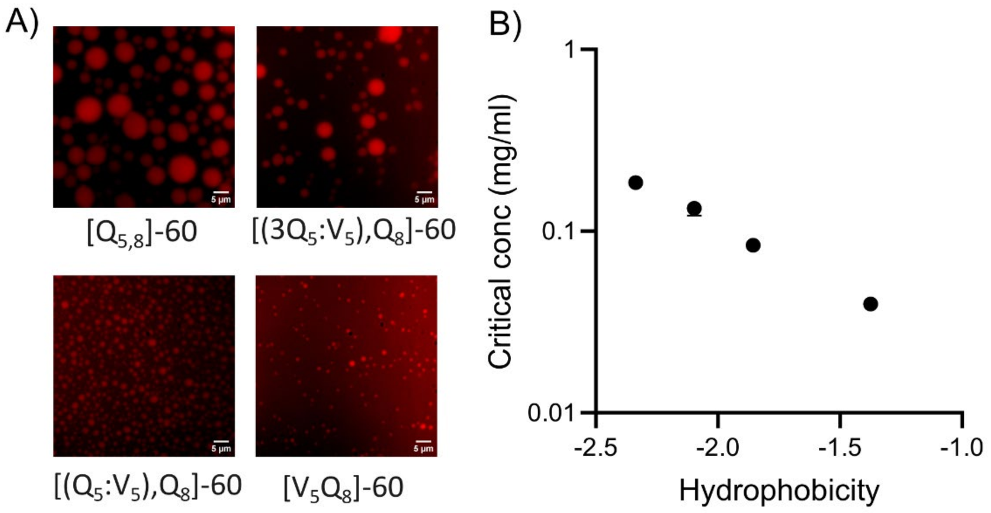


**Figure S6:** Characterization of the condensates formed by hydrophobic variations in $[Q_{5,8}]$-60. (A) Microscopic images of all the hydrophobic variants of $[Q_{5,8}]$-60; (B) Plot of the critical concentration required for phase separation v/s change in the hydrophobicity of the repeats.

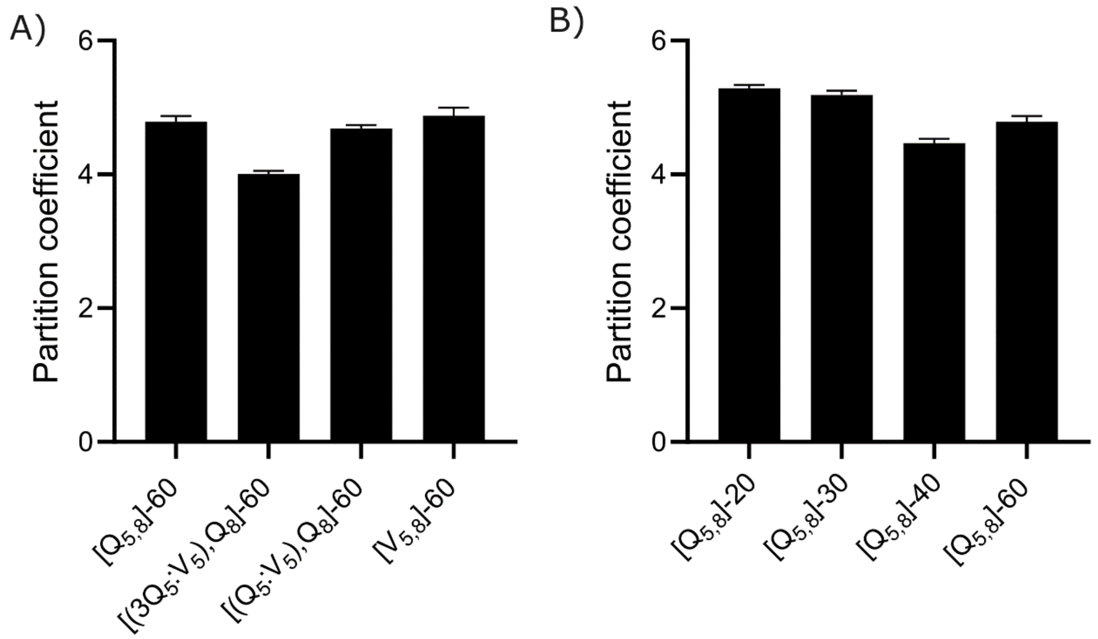


**Figure S7:** Partitioning coefficient of ruthenium dye is determined from ratio of fluorescence intensity of ruthenium dye inside and outside the droplets where A shows partition coefficient of the ruthenium dye in the hydrophobic variants of the $[Q_{5,8}]$-60 repeats and B shows partition coefficient of the dye into the $[Q_{5,8}]$-n repeats, where n varies from 20 to 60.

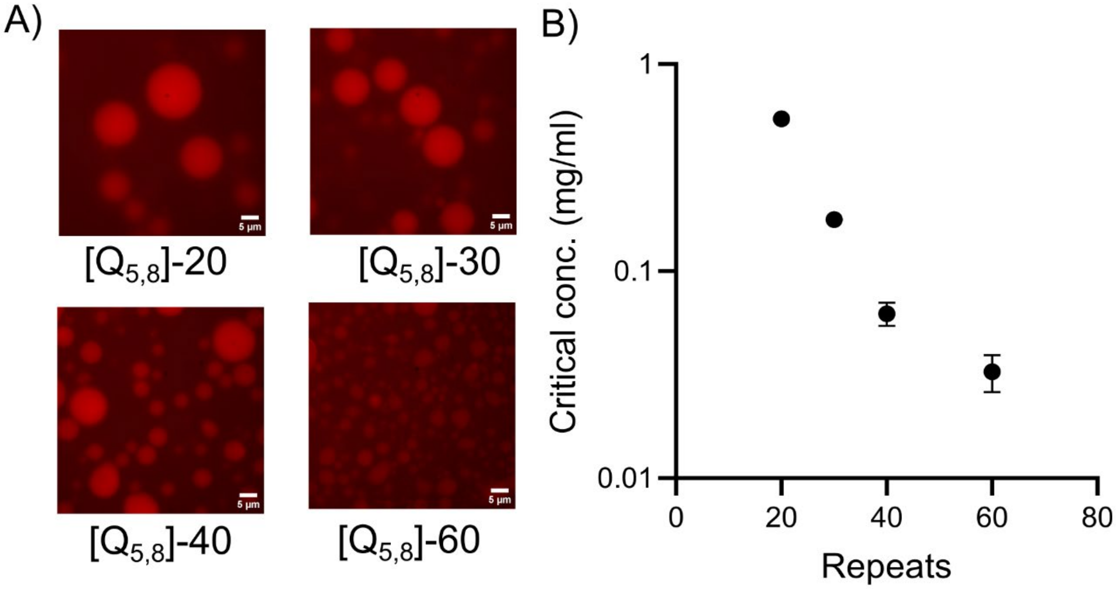


**Figure S8:** Characterization of the condensates formed by increasing peptide repeats. (A) Microscopic images of all the peptide repeats from 20 to 60 units; (B) Plot of the critical concentration required for phase separation v/s change in the number of the repeats.

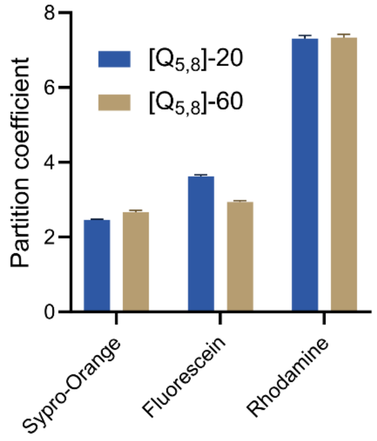


**Figure S9:** Partitioning of small molecule fluorescent dyes into condensates. The partitioning coefficient of the different hydrophobic dyes into [$Q_{5,8}$]-20 and [$Q_{5,8}$]-60 condensates calculated from intensity ratio in confocal microscopic images.

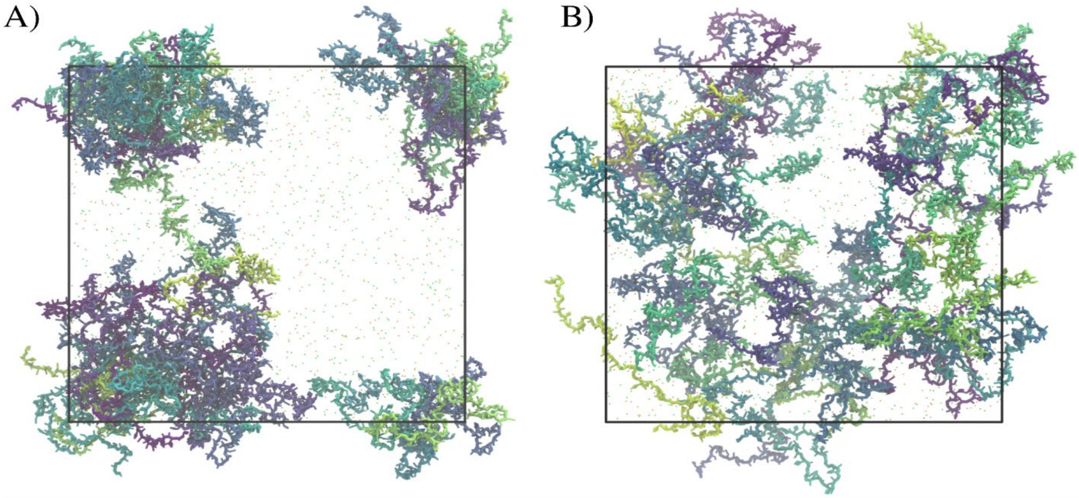

**Figure S10:** Optimizing water-protein interactions for reproducing biomolecular condensates of [$Q_{5,8}$]-20. A) Water-protein interactions increased by a factor of 1.04, where liquid-liquid phase separation has been achieved. B) Water-protein interactions scaled by a factor of 1.10, where proteins no longer interact with each other at all, and the system is homogeneous.

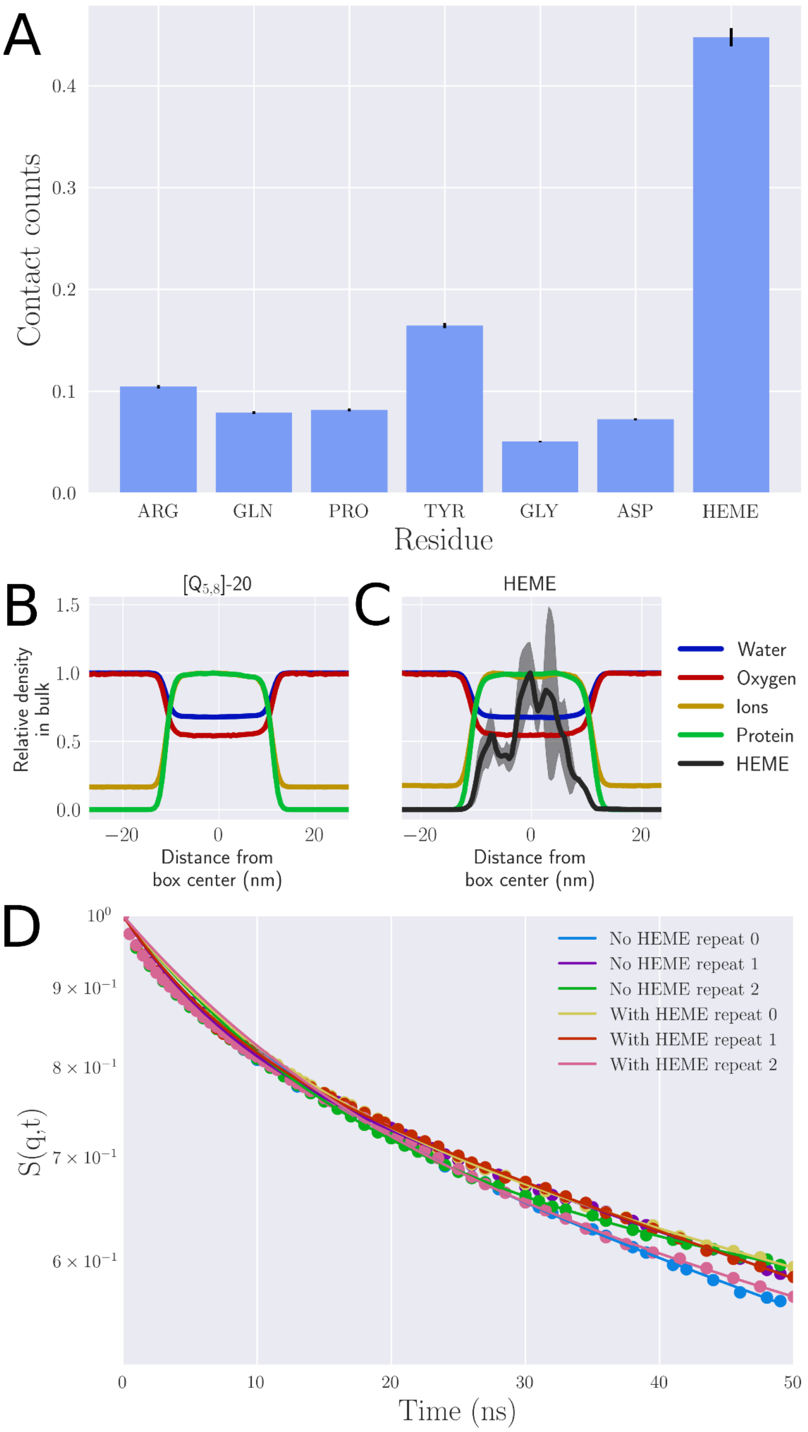


**Figure S11:** Partitioning of HEME into [$Q_{5,8}$]-20 condensates. A) Residue normalised contacts for oxygen against protein residues and HEME inside the condensate. B,C) System densities for the systems without (B) and with (C) HEME included in the simulation. The relative density of oxygen is identical in both systems (0.55). D) Incoherent scattering functions measured for proteins with and without HEME added to the system across three replicas for both conditions.

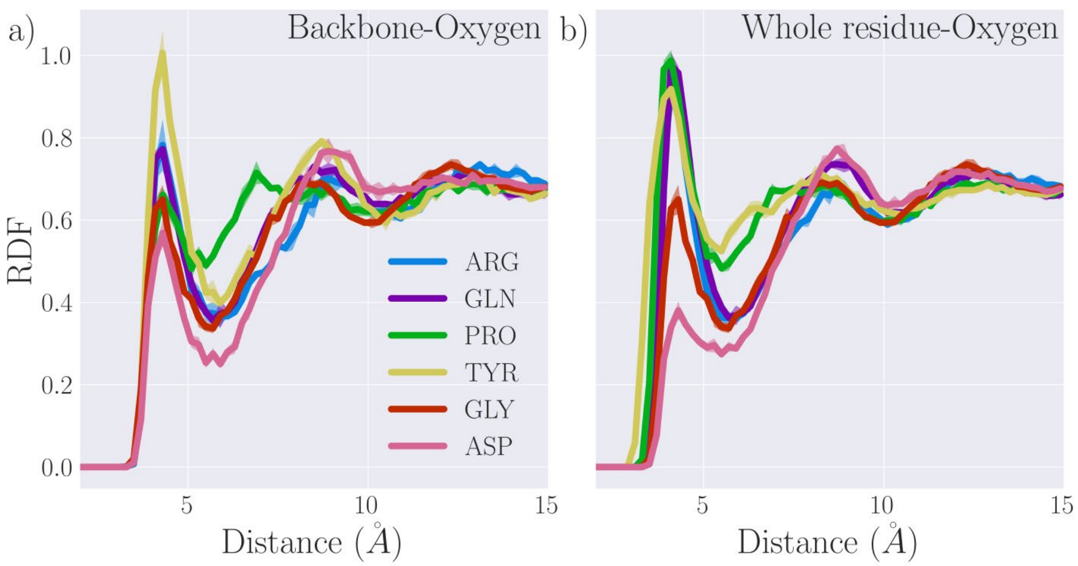


**Figure S12:** Comparison of residue-specific radial distribution function calculations in condensates. a) residue-specific RDFs as measured between protein backbone beads and oxygen molecules, reproduced from the main text. b) residue-specific RDFs measured between entire residues and oxygen molecules.

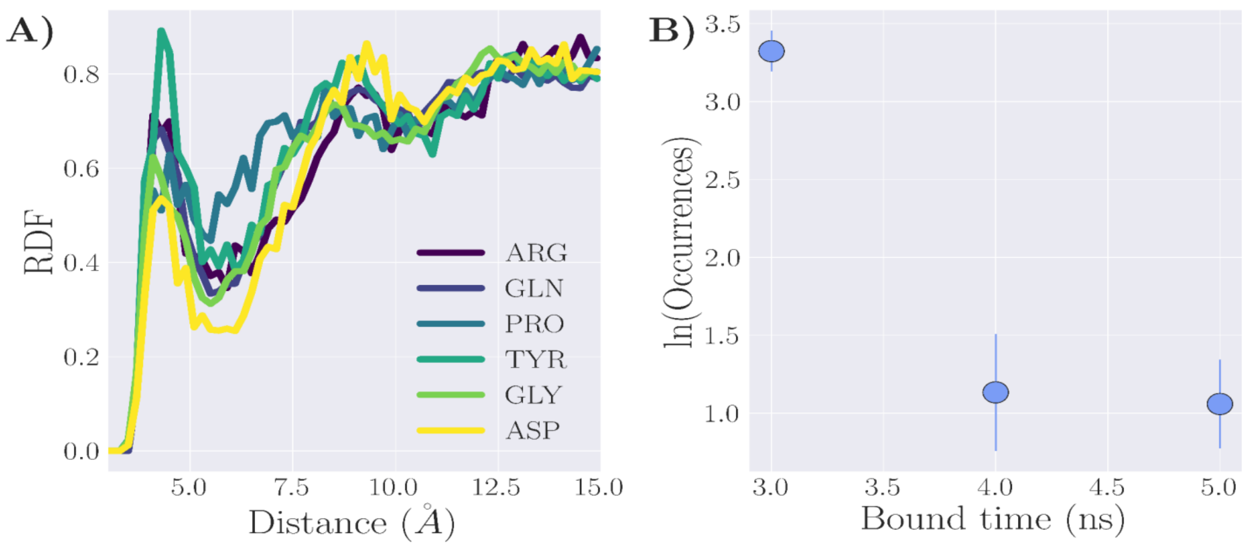


**Figure S13:** Interaction of oxygen with a protein in a dilute system. A) Residue-based radial distribution functions between residues of the [$Q_{5,8}$] repeat protein and oxygen. B) Prevalence of binding events between oxygen and the protein in the dilute system.

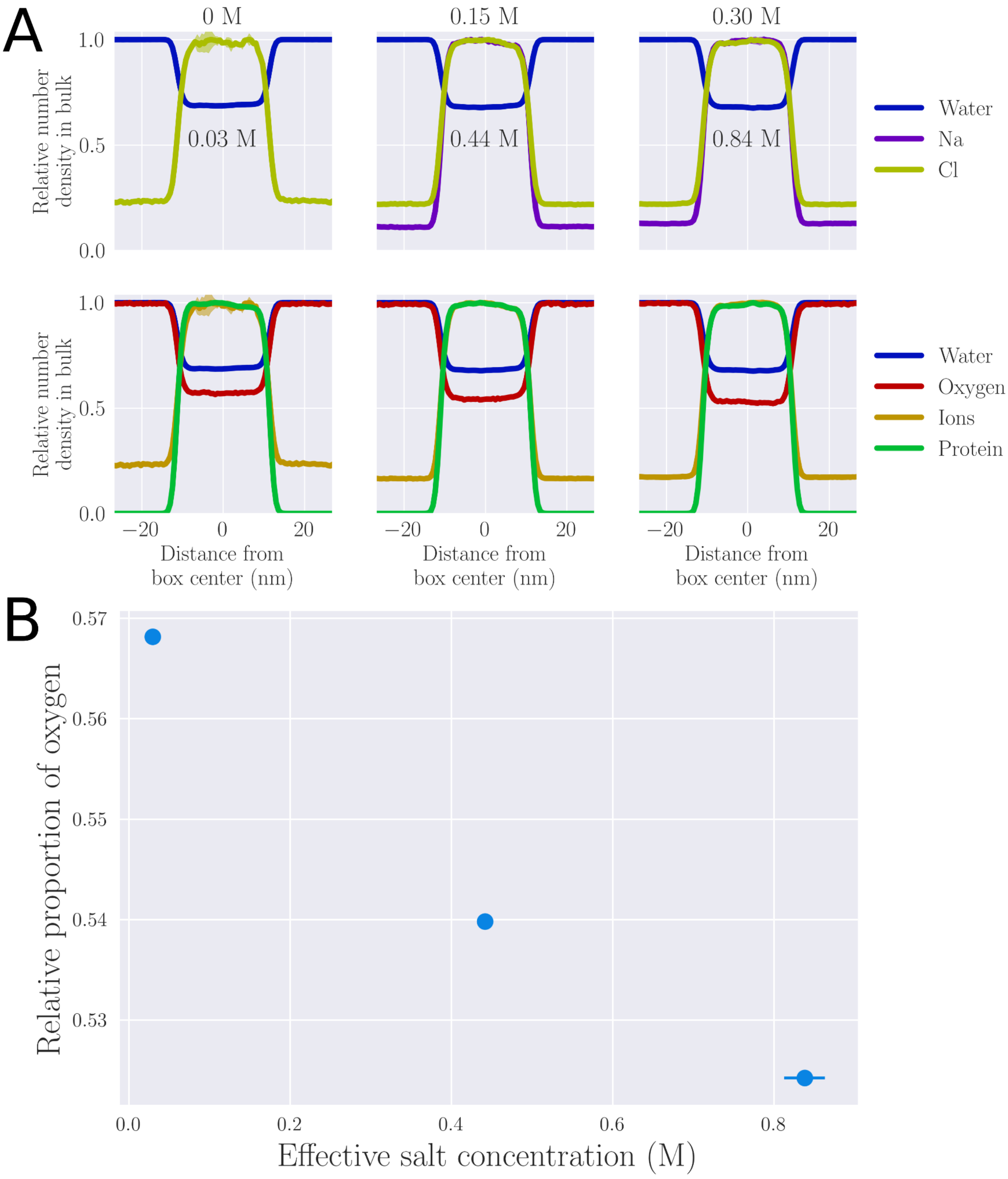


**Figure S14:** Effective salt concentration and oxygen partitioning. A) density profiles of system components showing the differential partitioning of the anion and cation, and the effective increase in salt concentration in the dense region of the simulated system. B) Summary of the effective salt concentration and the partitioning of oxygen into the dense phase of the system.

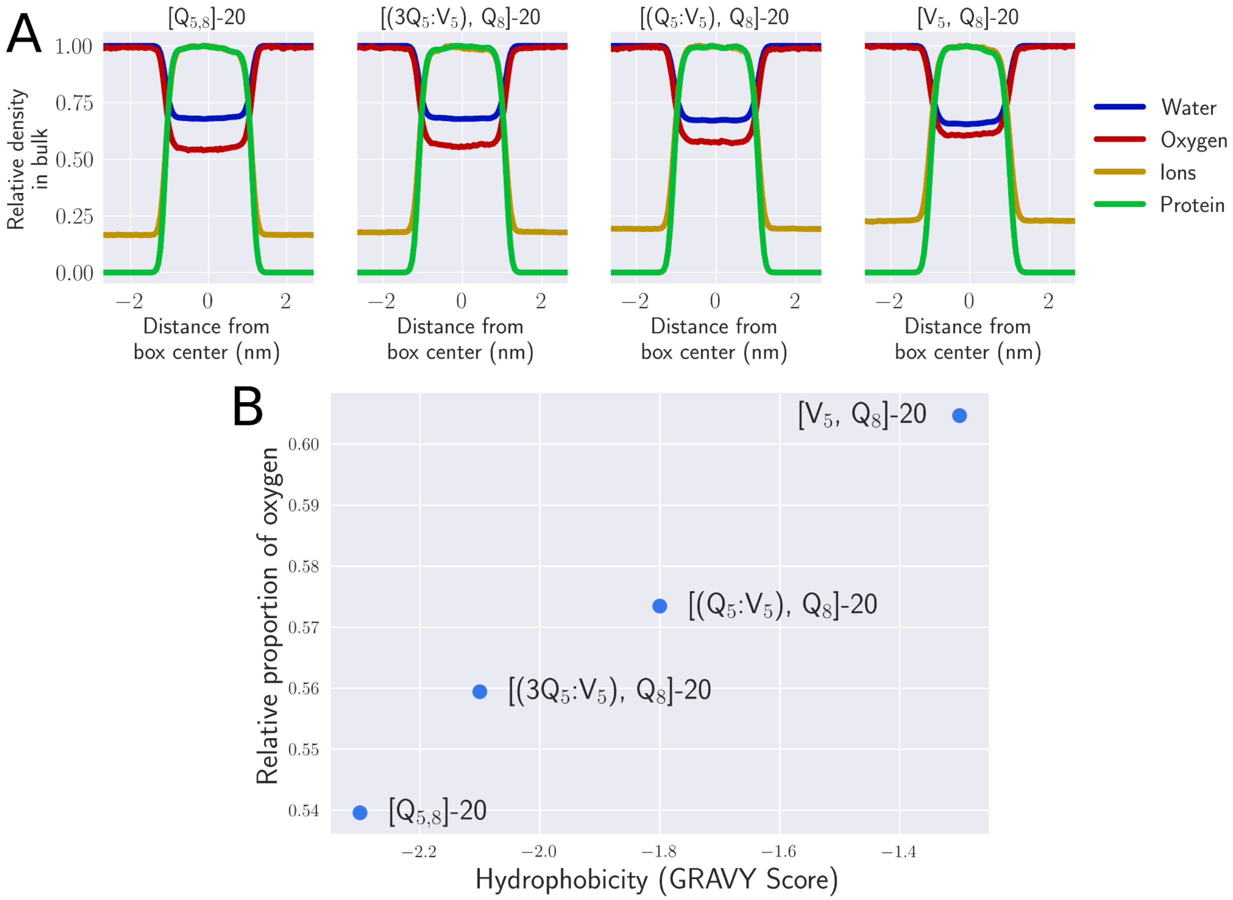


**Figure S15:** Effect of sequence in simulated condensates. A) Density profiles of the simulated constructs. B) Summary plot of the effect of sequence on oxygen partitioning

**Supporting Table**

**Table S1: Full amino acid sequence of the synthetic repeat peptides used in the study.**

| No. | Scaffold | Sequence |
|---|---|---|
| 1 | $[Q_{5,8}]$-20 | MGSSHHHHHHSSGLVPRGSHMSKGP-<br>[GRGDQPYQGRGDQPYQGRGDQPYQGRGDQPYQGRGD<br>QPYQGRGDQPYQGRGDQPYQGRGDQPYQGRGDQPYQG<br>RGDQPYQGRGDQPYQGRGDQPYQGRGDQPYQGRGDQP<br>YQGRGDQPYQGRGDQPYQGRGDQPYQGRGDQPYQGRG<br>DQPYQGRGDQPYQ]-GY |
| 2 | $[Q_{5,8}]$-30 | MGSSHHHHHHSSGLVPRGSHMSKGP-<br>[GRGDQPYQGRGDQPYQGRGDQPYQGRGDQPYQGRGD<br>QPYQGRGDQPYQGRGDQPYQGRGDQPYQGRGDQPYQG<br>RGDQPYQGRGDQPYQGRGDQPYQGRGDQPYQGRGDQP<br>YQGRGDQPYQGRGDQPYQGRGDQPYQGRGDQPYQGRG<br>DQPYQGRGDQPYQGRGDQPYQGRGDQPYQGRGDQPYQ<br>GRGDQPYQGRGDQPYQGRGDQPYQGRGDQPYQGRGDQ<br>PYQGRGDQPYQGRGDQPYQ]-GY |
| 3 | $[Q_{5,8}]$-40 | MGSSHHHHHHSSGLVPRGSHM-SKGP-<br>[GRGDQPYQGRGDQPYQGRGDQPYQGRGDQPYQGRGD<br>QPYQGRGDQPYQGRGDQPYQGRGDQPYQGRGDQPYQG<br>RGDQPYQGRGDQPYQGRGDQPYQGRGDQPYQGRGDQP<br>YQGRGDQPYQGRGDQPYQGRGDQPYQGRGDQPYQGRG<br>DQPYQGRGDQPYQGRGDQPYQGRGDQPYQGRGDQPYQ<br>GRGDQPYQGRGDQPYQGRGDQPYQGRGDQPYQGRGDQ<br>PYQGRGDQPYQGRGDQPYQGRGDQPYQGRGDQPYQGR<br>GDQPYQGRGDQPYQGRGDQPYQGRGDQPYQGRGDQPY<br>QGRGDQPYQGRGDQPYQGRGDQPYQ]-GY-GS |
| 4 | $[Q_{5,8}]$-60 | MGSSHHHHHHSSGLVPRGSHM-SKGP-<br>[GRGDQPYQGRGDQPYQGRGDQPYQGRGDQPYQGRGD<br>QPYQGRGDQPYQGRGDQPYQGRGDQPYQGRGDQPYQG<br>RGDQPYQGRGDQPYQGRGDQPYQGRGDQPYQGRGDQP<br>YQGRGDQPYQGRGDQPYQGRGDQPYQGRGDQPYQGRG<br>DQPYQGRGDQPYQGRGDQPYQGRGDQPYQGRGDQPYQ<br>GRGDQPYQGRGDQPYQGRGDQPYQGRGDQPYQGRGDQ<br>PYQGRGDQPYQGRGDQPYQGRGDQPYQGRGDQPYQGR |

| | | |
|---|---|---|
| | | GDQPYQGRGDQPYQGRGDQPYQGRGDQPYQGRGDQPY<br>QGRGDQPYQGRGDQPYQGRGDQPYQGRGDQPYQGRGD<br>QPYQGRGDQPYQGRGDQPYQGRGDQPYQGRGDQPYQG<br>RGDQPYQGRGDQPYQGRGDQPYQGRGDQPYQGRGDQP<br>YQGRGDQPYQGRGDQPYQGRGDQPYQGRGDQPYQGRG<br>DQPYQGRGDQPYQGRGDQPYQGRGDQPYQGRGDQPYQ]<br>-GY-GS |
| 5 | [3$Q_5$:$V_5$, Q8]-60 | MGSSHHHHHHSSGLVPRGS-HM-SKGP-<br>[GRGDQPYQGRGDQPYQGRGDQPYQGRGDVPYQGRGD<br>QPYQGRGDQPYQGRGDQPYQGRGDVPYQGRGDQPYQG<br>RGDQPYQGRGDQPYQGRGDVPYQGRGDQPYQGRGDQP<br>YQGRGDQPYQGRGDVPYQGRGDQPYQGRGDQPYQGRG<br>DQPYQGRGDVPYQGRGDQPYQGRGDQPYQGRGDQPYQ<br>GRGDVPYQGRGDQPYQGRGDQPYQGRGDQPYQGRGDV<br>PYQGRGDQPYQGRGDQPYQGRGDQPYQGRGDVPYQGR<br>GDQPYQGRGDQPYQGRGDQPYQGRGDVPYQGRGDQPY<br>QGRGDQPYQGRGDQPYQGRGDVPYQGRGDQPYQGRGD<br>QPYQGRGDQPYQGRGDVPYQGRGDQPYQGRGDQPYQG<br>RGDQPYQGRGDVPYQGRGDQPYQGRGDQPYQGRGDQP<br>YQGRGDVPYQGRGDQPYQGRGDQPYQGRGDQPYQGRG<br>DVPYQGRGDQPYQGRGDQPYQGRGDQPYQGRGDVPYQ]<br>-GY-GS |
| 6 | [$Q_5$:$V_5$, Q8]-60 | MGSSHHHHHHSSGLVPRGSHM-SKGP-<br>[GRGDQPYQGRGDVPYQGRGDQPYQGRGDVPYQGRGD<br>QPYQGRGDVPYQGRGDQPYQGRGDVPYQGRGDQPYQG<br>RGDVPYQGRGDQPYQGRGDVPYQGRGDQPYQGRGDVP<br>YQGRGDQPYQGRGDVPYQGRGDQPYQGRGDVPYQGRG<br>DQPYQGRGDVPYQGRGDQPYQGRGDVPYQGRGDQPYQ<br>GRGDVPYQGRGDQPYQGRGDVPYQGRGDQPYQGRGDV<br>PYQGRGDQPYQGRGDVPYQGRGDQPYQGRGDVPYQGR<br>GDQPYQGRGDVPYQGRGDQPYQGRGDVPYQGRGDQPY<br>QGRGDVPYQGRGDQPYQGRGDVPYQGRGDQPYQGRGD<br>VPYQGRGDQPYQGRGDVPYQGRGDQPYQGRGDVPYQG<br>RGDQPYQGRGDVPYQGRGDQPYQGRGDVPYQGRGDQP<br>YQGRGDVPYQGRGDQPYQGRGDVPYQGRGDQPYQGRG<br>DVPYQGRGDQPYQGRGDVPYQGRGDQPYQGRGDVPYQ]<br>-GY-GS |
| 7 | [$V_5$, Q8]-60 | MGSSHHHHHHSSGLVPRGS-HM-SKGP-<br>[GRGDVPYQGRGDVPYQGRGDVPYQGRGDVPYQGRGD |

| | | |
|---|---|---|
| | | VPYQGRGDVPYQGRGDVPYQGRGDVPYQGRGDVPYQGRGDVPYQGRGDVPYQGRGDVPYQGRGDVPYQGRGDVPYQGRGDVPYQGRGDVPYQGRGDVPYQGRGDVPYQGRGDVPYQGRGDVPYQGRGDVPYQGRGDVPYQGRGDVPYQGRGDVPYQGRGDVPYQGRGDVPYQGRGDVPYQGRGDVPYQGRGDVPYQGRGDVPYQGRGDVPYQGRGDVPYQGRGDVPYQGRGDVPYQGRGDVPYQGRGDVPYQGRGDVPYQGRGDVPYQGRGDVPYQGRGDVPYQGRGDVPYQGRGDVPYQGRGDVPYQGRGDVPYQGRGDVPYQGRGDVPYQGRGDVPYQGRGDVPYQGRGDVPYQGRGDVPYQGRGDVPYQGRGDVPYQGRGDVPYQGRGDVPYQGRGDVPYQGRGDVPYQGRGDVPYQGRGDVPYQGRGDVPYQGRGDVPYQGRGDVPYQGRGDVPYQGRGDVPYQGRGDVPYQGRGDVPYQGRGDVPYQ]-GY-GS |
| **Note: 6X His tag coded in red color.**<br><br>**valine substitutions are coded in blue.** | | |
